# Epigenomic Diversity of Cortical Projection Neurons in the Mouse Brain

**DOI:** 10.1101/2020.04.01.019612

**Authors:** Zhuzhu Zhang, Jingtian Zhou, Pengcheng Tan, Yan Pang, Angeline Rivkin, Megan A. Kirchgessner, Elora Williams, Cheng-Ta Lee, Hanqing Liu, Alexis D. Franklin, Paula Assakura Miyazaki, Anna Bartlett, Andrew Aldridge, Minh Vu, Lara Boggeman, Conor Fitzpatrick, Joseph R. Nery, Rosa G. Castanon, Mohammad Rashid, Matthew Jacobs, Tony Ito, Bertha Dominguez, Sheng-Yong Niu, Jared B. Smith, Carolyn O’Connor, Kuo-Fen Lee, Xin Jin, Eran A. Mukamel, M. Margarita Behrens, Joseph R. Ecker, Edward M. Callaway

**Affiliations:** Genomic Analysis Laboratory, The Salk Institute for Biological Studies, La Jolla, CA 92037; Bioinformatics and Systems Biology Program, University of California San Diego, La Jolla, CA 92093; School of Pharmaceutical Sciences, Tsinghua University, Beijing, China, 100084; Systems Neurobiology Laboratories, The Salk Institute for Biological Studies, La Jolla, CA 92037; Neurosciences Graduate Program, University of California, San Diego, La Jolla, CA 92093; Molecular Neurobiology Laboratory, The Salk Institute for Biological Studies, La Jolla, CA 92037; Peptide Biology Laboratories, The Salk Institute for Biological Studies, La Jolla, CA 92037; Division of Biological Sciences, University of California San Diego, La Jolla, CA 92093; Flow Cytometry Core Facility, The Salk Institute for Biological Studies, La Jolla, CA 92037; Department of Cognitive Science, University of California, San Diego, La Jolla, CA 92037; Computational Neurobiology Laboratory, The Salk Institute for Biological Studies, La Jolla, CA 92037; Howard Hughes Medical Institute, The Salk Institute for Biological Studies, La Jolla, CA 92037

## Abstract

Neuronal cell types are classically defined by their molecular properties, anatomy, and functions. While recent advances in single-cell genomics have led to high-resolution molecular characterization of cell type diversity in the brain, neuronal cell types are often studied out of the context of their anatomical properties. To better understand the relationship between molecular and anatomical features defining cortical neurons, we combined retrograde labeling with single-nucleus DNA methylation sequencing to link epigenomic properties of cell types to neuronal projections. We examined 11,827 single neocortical neurons from 63 cortico-cortical (CC) and cortico-subcortical long-distance projections. Our results revealed unique epigenetic signatures of projection neurons that correspond to their laminar and regional location and projection patterns. Based on their epigenomes, intra-telencephalic (IT) cells projecting to different cortical targets could be further distinguished, and some layer 5 neurons projecting to extra-telencephalic targets (L5-ET) formed separate subclusters that aligned with their axonal projections. Such separation varied between cortical areas, suggesting area-specific differences in L5-ET subtypes, which were further validated by anatomical studies. Interestingly, a population of CC projection neurons clustered with L5-ET rather than IT neurons, suggesting a population of L5-ET cortical neurons projecting to both targets (L5-ET+CC). We verified the existence of these neurons by labeling the axon terminals of CC projection neurons and observed clear labeling in ET targets including thalamus, superior colliculus, and pons. These findings highlight the power of single-cell epigenomic approaches to connect the molecular properties of neurons with their anatomical and projection properties.

## Main Text

The mammalian brain is a complex system consisting of multiple types of neurons with diverse morphology, physiology, connections, gene expression, and epigenetic modifications. Identifying brain cell types and how they interact is critical to understanding the neural mechanisms that underlie brain function. During the last decade, these efforts have been facilitated by the advent of molecular, genetic and viral tools for allowing genetic access and manipulation of specific cell types^1,2^. Available evidence suggests, however, that there are far more cell types than can presently be accessed genetically. Moreover, the correspondence between molecular cell types and neuronal populations defined by connectivity are largely unknown.

Single-cell technologies deconvolve mammalian brains into molecularly defined cell clusters corresponding to putative neuron types^3^. Among these technologies, single nucleus methylation sequencing (snmC-Seq) applied to neurons has the unique ability to allow identification of potential regulatory elements and a prediction of gene expression in the same cells. This is because methylation at non-CG (CH; H= A, T, C) dinucleotides (mCH) of the gene body is inversely correlated with RNA expression, and methylation at both CG dinucleotides (mCG) and CH dinucleotides can be used to identify gene regulatory elements associated with gene expression^4–6^. Furthermore, CH methylation accumulates and CG methylation reconfigures during cortical synaptic development, suggesting possible links between epigenetics and connectivity^7,8^.

Previous single-cell analyses have revealed transcriptomic clusters and linked them to neuron types with different projection patterns in a few particular brain regions^9–12^. For the cerebral cortex, the most prominent molecular distinction related to projection targets is the separation of cortical neurons into distinct and apparently non-overlapping IT and L5-ET (also called pyramidal tract, PT) groups. In some cases L5-ET cells have been further divided based on both gene expression and corresponding axon projections^9^. While the separation of L5-IT and ET neurons appears to be conserved across cortical areas^13^ and species^14^, a systematic analysis of the relationships between a larger set of projection targets and molecular identities across multiple cortical areas has not been conducted. To what extent cortical projection neuron types can be further distinguished or divided by incorporating anatomical information with molecular analyses, and whether these cell types and correspondences are conserved across cortical areas is unclear. Ultimately, the use of methods that can classify cell types and predict regulatory elements, such as snmC-seq, will be critical to understanding cell type and/or projection type specific regulatory mechanisms.

To address these questions we developed Epi-Retro-Seq, which applies snmC-Seq^15^ to neurons dissected from cortical source regions which were labeled based on their long distance projections to specific cortical and subcortical targets. We analyzed the methylomes of 11,827 single neurons from eight cortical areas projecting to ten target regions. This dataset enabled us to quantify the epigenetic differences between cortical projection neurons, to identify specific genes and regulatory elements in projection neurons, to study the relationships between cortical projection neurons and molecular cell types, and to identify a neuron type making projections to both cortical and ET targets.

## Results

### Epi-Retro-Seq of 63 cortical projections

To obtain a comprehensive view of the molecular diversity among cortical projection neurons we performed Epi-Retro-Seq, which combines retrograde tracing with epigenomic profiling. We characterized projection neurons from eight cortical areas (“source”) spanning the anterior-to-posterior extent of the mouse cortex that project to ten cortical or subcortical regions (“target”) (Fig. 1a), covering overall 26 CC projections and 37 cortico-subcortical projections (Supplementary Table 1). In Epi-Retro-Seq, the retrograde viral tracer rAAV2-retro-Cre is injected in the target region in an INTACT mouse^4^, turning on Cre-dependent nuclear-GFP expression in neurons that project to the injected target, throughout the mouse brain. The brain is then sectioned into eighteen 600-micron coronal slices, and the source regions of interest are dissected from each slice (see Methods). Nuclei are sampled from at least 4 mice (2 male and 2 female) for each projection target (except AI→pons - 2 male mice only). Nuclei from each of the dissected source regions are prepared, from which GFP^+^/NeuN^+^ nuclei (the GFP-labeled projection neurons) are isolated as single nuclei using fluorescence activated nuclei sorting (FANS) and assayed using snmC-Seq2^15^ to profile their genome-wide DNA methylation signatures. The ten injected target regions include four cortical areas [the primary motor cortex (MOp), primary somatosensory cortex (SSp), anterior cingulate area (ACA), and primary visual cortex (VISp)], and six major subcortical structures [the striatum (STR), thalamus (TH), superior colliculus (SC), ventral tegmental area and substantia nigra (VTA+SN), pons, and medulla (MY)]. Each of the eight source cortical regions [MOp, SSp, ACA, agranular insular cortex (AI), retrosplenial cortex (RSP), auditory cortex (AUDp+AUDd+AUDv), posterior parietal cortex (PTLp), and visual cortex (VISp+VISpm+VISl+VISli)] were hand dissected from one or two coronal slices following the Allen Mouse Common Coordinate Framework (CCF), Reference Atlas, Version 3 (2015) (Extended Data Fig. 1).

**Fig. 1.**
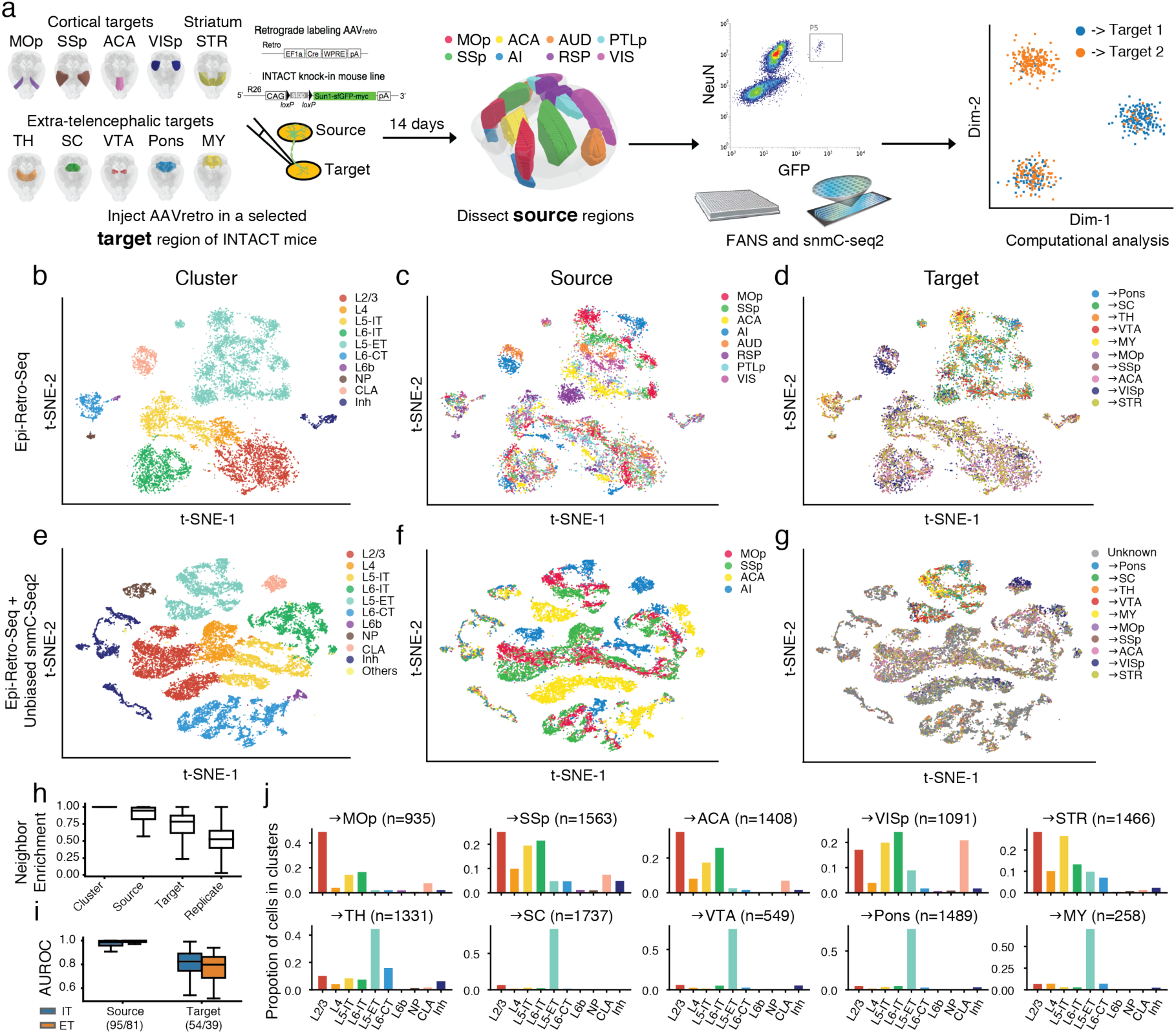
The epigenomic landscape of cortical projection neurons. **a**, Schematics of Epi-Retro-Seq workflow that retrogradely labels and epigenetically profiles single projection neurons. The retrograde tracer rAAV2-retro-cre was injected in one of the ten target regions (primary motor cortex (MOp), primary somatosensory cortex (SSp), anterior cingulate cortex (ACA), primary visual cortex (VISp), striatum (STR), thalamus (TH), superior colliculus (SC), the ventral tegmental area (VTA) & substantia nigra (SNr), Pons, or medulla (MY)) in INTACT knock-in mice. Therefore, nuclei of neurons that projected to the injected target were labeled with cre-dependent nuclear GFP. Source regions of interest (MOp, SSp, ACA, agranular insular cortex (AI), auditory cortex (AUD), retrosplenial cortex (RSP), posterior parietal cortex (PTLp), or visual cortex (VIS)) were dissected 14 days after the injection, from which nuclei were prepared and single GFP^+^/NeuN^+^ nuclei were isolated using fluorescence activated nuclei sorting (FANS) followed by snmC-seq2 and computational analysis. Brain diagrams were derived from the Allen Mouse Brain Reference Atlas (version 3 (2015)). **b-d**, Two-dimensional t-distributed stochastic neighbor embedding (t-SNE) of 11,827 cortical neuron nuclei based on CH methylation (mCH) levels in 100 kb genomic bins, colored by cluster (**b**), the source region of neurons (**c**), or their projection target (**d**). Cortical neurons were better separated by their source regions than projection targets within each major cell type cluster. **e-g**, Integrative clustering of Epi-Retro-Seq and unbiased snmC-seq2 (without enrichment of projections) of neurons from MOp, SSp, ACA and AI (n=21,966), colored by cluster (**e**), source region (**f**), and projection targets in Epi-Retro-Seq (**g**). **h**, Neighbor enrichment scores of cells (n=11,827) categorized by cluster, source, target, and replicate. **i**, AUROC of source pairs and target pairs computed for IT (blue) and ET (orange) neurons based on gene body mCH. Sample sizes are shown in x-axis ticklabels. **j**, The distribution across cell clusters of neurons that projected to each IT (top row) or ET (bottom row) target. The elements of all boxplots are defined as: center line, median; box limits, first and third quartiles; whiskers, 1.5× interquartile range. IT, intra-telencephalic; ET, extra-telencephalic; NP, near-projecting; CT, corticothalamic; Inh, inhibitory; CLA, claustrum; Others, cell clusters detected in unbiased snmC-seq2 but not in Epi-Retro-Seq.

### Methylation landscape of cortical projection neurons

We assayed approximately 384 nuclei from each projection (except the MOp→SSp projection from which 768 nuclei were assayed). After removing the low-quality cells, potential doublets, and glial cells (possibly due to false NeuN positives in FANS), we obtained high-quality single methylomes for 11,827 cortical projection neurons (Extended Data Fig. 2). The level of CH methylation in each single nucleus was computed across the genome using 100 kb genomic bins and used to perform unsupervised clustering of the projection neurons. Overall, the cortical projection neuron clusters were annotated into 10 major cell types (Fig. 1b) based on the reduced levels of gene body mCH, a proxy for gene expression, of known marker genes (Extended Data Fig. 2f). It should be noted that 361 neurons (3.05%) fell into the inhibitory neuron cluster, likely representing false-positives possibly, due to either labeling of neurons by AAV that leaked into cortical areas above subcortical injection sites (mostly from areas above TH injections), or insufficient gating stringency during FANS, allowing inclusion of GFP-negative nuclei. This low error rate allows a rough estimate of the likely erroneous contributions from other cell types. Within each cell type cluster, excitatory neurons but not inhibitory neurons from different cortical regions were further separated from each other (Fig. 1c), demonstrating that such separations in excitatory neuron clusters were not due to technical effects but instead represented the distinct spatial DNA methylation patterns in cortical projection neurons. As can be seen from the t-SNE visualization (Fig. 1d), neurons projecting to different target regions were more similar within each cluster than neurons from different source regions, indicating that they shared a more similar DNA methylation landscape. Neighbor enrichment scores were used to quantify the variations of DNA methylation that originated from different cell types, cortical spatial regions, and projection targets (see Methods). Neurons from the same cluster occupied highly similar regions in the dimension reduction space (neighbor enrichment score was near 1). Scores were also high for comparisons across neurons from the same source, followed by projections to the same target. Scores were near chance for biological replicates (Fig. 1h).

Next, we integrated our data with the single-nuclei methylation data that were dissected and sorted from some of the same cortical regions but without enrichment of specific projections (Liu et al., companion paper #9). We observed a close agreement of the major cell types (Fig. 1e) and source regions (Fig. 1f) between these two datasets. Given the increased number of cells, different source regions became better demarcated on t-SNE (Fig. 1f). Compared with unbiased snmC-seq2 profiling, Epi-Retro-seq dataset also contains information about the neuronal projection targets revealed by retrograde tracing (Fig. 1g). This enabled enrichment of rare types of projection neurons and analysis of the methylation patterns of neurons projecting to different brain regions.

Although neurons projecting to different target regions were not completely separated on t-SNE, we observed an explicit enrichment of CC and cortico-striatal projection neurons in IT clusters (L2/3, L4, L5-IT, L6-IT, and Claustrum (CLA)), separated from neurons that project to the remaining structures outside the telencephalon which were categorized as L5-ET neurons (Fig. 1j, Extended Data Fig. 3) As expected, many cortico-thalamic projecting neurons were also found in the L6-CT cluster (Fig. 1j, Extended Data Fig. 3). These enrichment patterns are consistent with our knowledge about laminar enrichment of the projection neurons, which reflects the high quality of our retrogradely labeled single-nuclei methylation dataset.

To further quantify methylation differences between neurons from different source regions or projecting to different target regions, we made comparisons across source pairs or target pairs. For each pair of interest, area under the curve of receiver operating characteristic (AUROC) was calculated to score the level of separation between the two groups of projection neurons. Specifically, a logistic regression model was trained using normalized gene body mCH as features to predict which group a cell belongs to. By training the model in one biological replicate and testing on the other, the performance was measured by AUROC. By comparing each pair of sources or targets, we found that most neurons dissected from different source regions could be separated with AUROC > 0.9 (Fig. 1i). Most of the neurons projecting to different target regions were also separable by mCH in this supervised setting (Fig. 1i), although they were closely mixed in the unsupervised embeddings (Fig. 1d). These findings indicate that nearly all of the different types of projection neurons that were profiled have differences in their epigenomes.

### Epigenetic diversity of IT neurons projecting to different cortical targets

As described above, assessment of the entire Epi-Retro-Seq dataset revealed clear and expected differences in the neuron clusters occupied by neurons projecting to IT versus ET targets, and these differences were conserved across source areas. However, neurons projecting to different IT or ET targets did not uniquely separate into distinct clusters when analyzed at the level of the entire cell population. Nevertheless, we were able to detect projection-dependent quantitative differences in the levels of DNA methylation. Further analyses of these quantitative differences, described below, allowed assessment of possible organizational principles that might exist in the relationships between DNA methylation, projections targets, and sources, including both areal and laminar sources.

In total, 42.6% of the cortical projection neurons profiled in our Epi-Retro-Seq data were identified as IT, and annotated according to their presumptive cortical layers (Fig. 1b). We next aimed to disentangle the contribution of the cortical area in which cell bodies were located versus their cortical projection targets, to the variation of their DNA methylation profiles. We focused on 26 CC projections from 8 cortical areas to 4 different cortical targets. AUROC scores were used to evaluate epigenetic relationships between cortical neurons projecting to different cortical targets. All possible pairs of 4 cortical targets were assessed for each of the 8 sources to generate 29 AUROC scores, organized according to projection target pairs (Fig. 2a, Extended Data Fig. 4a, c). Significant differences were observed between projection target pairs when assessed across source areas (p=6.8e-3, Kruskal-Wallis test), but not between cortical areas when assessed across target pairs (p=0.3, Kruskal-Wallis test). Among the six projection target pairs examined, neurons projecting to MOp versus ACA were overall most distinguishable (average AUROC = 0.902), followed by neurons projecting to ACA versus VISp (average AUROC = 0.887), while neurons that project to SSp versus ACA were the least separable (average AUROC = 0.693) (Fig. 2a). In addition, for each target pair, the performance of the predictive model varied among neurons from different source cortical regions (Fig. 2a, Extended Data Fig. 4a, c).

**Fig. 2.**
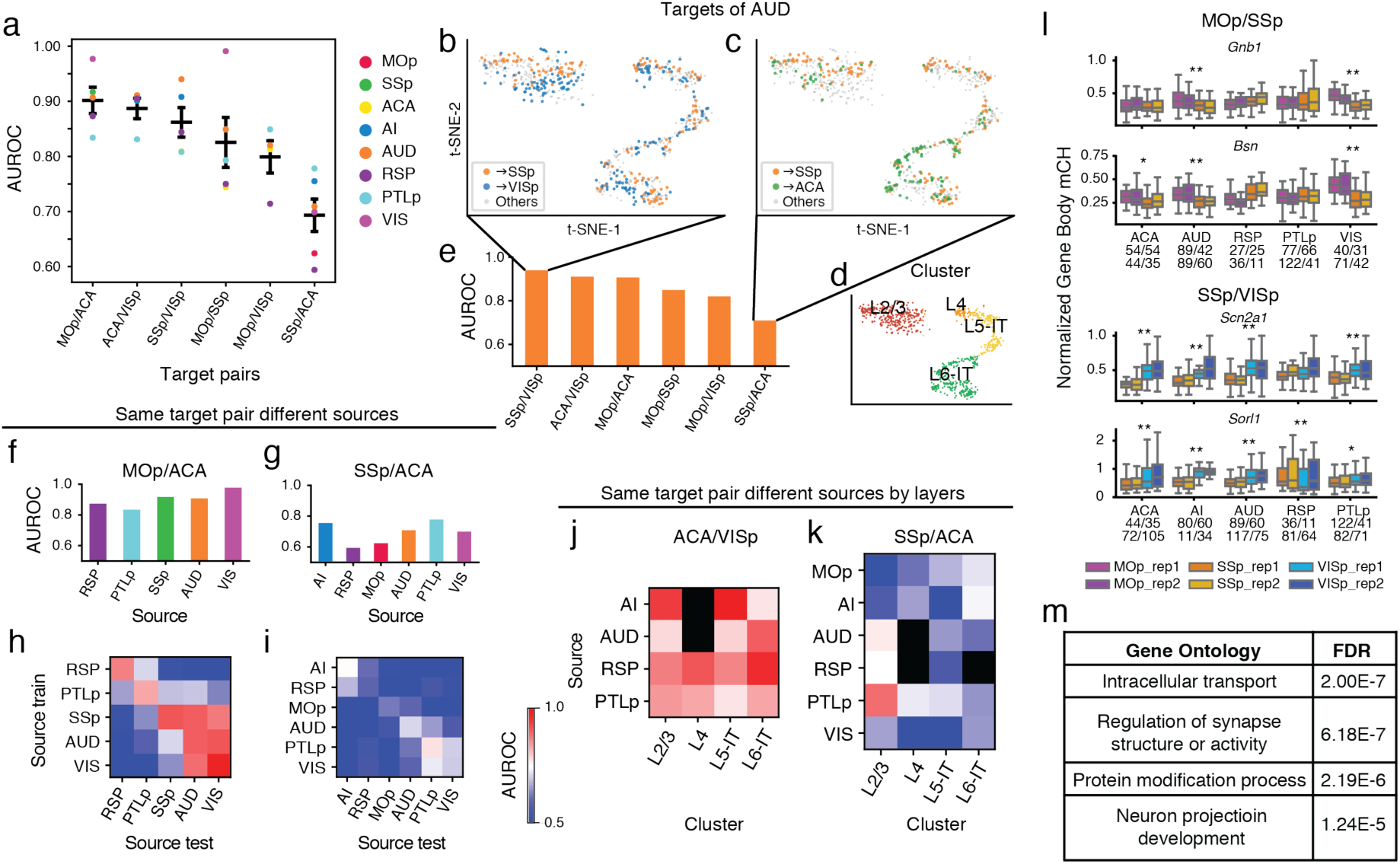
Epigenetic differences between IT neurons projecting to different targets. **a**, AUROC from the prediction model constructed to distinguish cortical neurons projecting to one cortical target versus another was used to measure the epigenetic variation between different cortical IT neurons. A significant variation of AUROC among different projection target pairs was observed. **b-e**, Upon examining AUD IT neurons (n=737) that project to different cortical targets, AUD→SSp neurons and AUD→VISp neurons were biased toward different locations within each layer-annotated cluster (**d**) on the t-SNE plot using mCH levels in gene bodies (**b**), while AUD→SSp neurons and AUD→ACA neurons were more intermingled (**c**). The differential levels of separation on t-SNE corresponded to the high AUROC between AUD→SSp versus AUD→VISp neurons, and low AUROC between AUD→SSp versus AUD→ACA neurons (**e**). **f, g**, The AUROC for comparisons between →MOp versus →ACA neurons from different source regions varied between 0.834 and 0.977 (**f**), while the AUROC for comparisons between →SSp versus →ACA neurons from different source regions varied between 0.594 and 0.778 (**g**), indicating overall higher levels of distinguishability between →MOp versus →ACA neurons, than between →SSp versus →ACA neurons. **h, i**, Heatmaps of AUROC from prediction models that were trained on one source region (row) and tested on another source region (column) to distinguish between neurons projecting to →MOp versus →ACA (**h**), or between →SSp versus →ACA neurons (**i**). **j, k**, Heatmaps of AUROC from prediction models that were trained and tested on neurons from each cortical layer (column) in each source region (row), to distinguish between →ACA versus →VISp neurons (**j**), or between →SSp versus →ACA neurons (**k**). **l**, Boxplots of example genes that were differentially methylated at CH sites (CH-DMGs) between →MOp versus →SSp neurons (top), or between →SSp versus →VISp neurons (bottom). The sample sizes are shown as ticklabels of x-axis. ** represents false discovery rate (FDR)<0.01 and * represents FDR<0.1. **m**, Gene ontology (GO) enrichment of 1,830 CH-DMGs between cortical neurons projecting to different cortical targets. The elements of all boxplots are defined as: center line, median; box limits, first and third quartiles; whiskers, 1.5× interquartile range. Center lines and error bars in (a) represent the means and standard errors of the means.

Together, these analyses suggest that epigenetic differences between CC projection neurons depend on a combination of both the specific targets to which neurons project and the source region where the neurons reside. For example, we further evaluated the variability of mCH profiles among AUD IT neurons projecting to different targets and found that AUD→SSp neurons were better separated from AUD→VISp neurons (AUROC = 0.94; Fig.2b, e) than from AUD→ACA neurons (AUROC = 0.709; Fig. 2c, e). t-SNE plots color-coded according to these same projection comparisons (Fig. 2b, c) or according to annotated layers (Fig. 2d) allow visualization of the extent to which these neurons differ. In addition to the apparent greater separability of AUD→SSP versus AUD→VISp than AUD→SSP versus AUD→ACA neurons, it can be seen that the distinctions between these projections did not stem from different distributions across layers (Fig. 2d). This demonstrates that the level of epigenetic differences between AUD IT neurons varies depending on their projection targets. On the other hand, when comparing neurons from different sources projecting to the same target pair, we observed different levels of distinguishability in our models. For example, while MOp-projecting versus ACA-projecting neurons were more distinguishable (i.e. higher AUROC scores) than SSp-projecting versus ACA-projecting neurons, we observed variation of the AUROC scores across different source regions for both target pairs (Fig. 2f, g).

To further validate that the differences in separability across regions resulted from biological differences rather than limited sample sizes for some regions, we trained our predictive model between two targets using neurons from one source region and then tested the performance of the model on another source region. These analyses also allowed evaluation of whether the same epigenetic differences that distinguished target pairs for one source area might be conserved across source areas. As expected, the performances of the cross-source-region models in distinguishing two projection targets were usually less than the same-source-region models (Fig. 2h, i, Extended Data Fig. 4b, d). Nevertheless, many target pairs that were distinguishable for the within-source models were also distinguishable with the cross-source models (Fig. 2h, i, Extended Data Fig. 4b, d), indicating conservation of target pair epigenetic differences across sources. Interestingly, the performance of models trained on any particular region varied in their ability to predict projections from other regions. For example, the model trained on data from AUD performed better in distinguishing VIS→MOp versus VIS→ACA neurons than the models trained on RSP, PTLp, or SSp (Fig. 2h). This suggests that AUD and VIS neurons are more similar to each other in the molecular markers that distinguish neurons projecting to MOp versus ACA than other cortical areas. These results indicate that cortical regions might form different groups with shared correlations between molecular markers and projection targets.

In addition, the level of distinguishability between two cortical targets appeared to be similar across layers (Fig. 2j, Extended Data Fig. 5a, b). By training and testing the predictive models in each layer separately, we observed higher distinguishability between ACA-projecting versus VISp-projecting neurons across all layers than between SSp-projecting versus ACA-projecting in all layers in almost all source regions (Fig. 2j, k). We further tested if cross-layer-trained models could distinguish the projection targets (see Methods), and observed that the performance was generally comparable to within-layer models (Extended Data Fig. 5c, d). These results suggest that there may be shared epigenetic signatures across layers that contribute to correlations with the projection targets.

To better understand the biology underlying the epigenetic signatures that distinguish different cortical IT projection neurons, we identified differentially methylated genes at CH sites (CH-DMGs) between different pairs of CC projection neurons in each source region using hierarchical linear models. In total, 1830 CH-DMGs were identified (Supplementary Table 3), among which 1,623 (88.7%) were statistically significant in only one source region, and 207 (11.3%) were differentially methylated in more than one source region (some examples shown in Fig. 2l). That the vast majority of CH-DMGs were unique to one source region, suggests that different genes may participate in defining projections from different source regions. Gene ontology (GO) enrichment analysis revealed that CH-DMGs were enriched for genes that participate in intracellular transport, regulation of synapse structure, etc. (Fig. 2m), all relevant for influencing neuronal projections. For example, Bassoon (*Bsn*) is differentially methylated between MOp-projecting and SSp-projecting neurons in ACA, AUD, and VIS (Fig. 2l). It encodes a presynaptic cytomatrix protein expressed primarily in neurons, and is essential in regulation of neurotransmitter release^16^. *Scn2a1* encodes a voltage dependent sodium channel protein and is differentially methylated between SSp-projecting and VISp-projecting neurons in ACA, AI, AUD, and PTLp (Fig. 2l). This channel regulates neuronal excitability and variants are associated with autism and seizure disorders^17^.

### Epigenetically distinct subpopulations of L5-ET neurons

In our Epi-Retro-Seq data, 5 out of the 10 profiled projection targets are ET. In particular, L5-ET neurons are the most abundant cell population in our datasets (4,176 (35.3%) single neurons), and are 6.3 fold enriched in Epi-Retro-Seq compared to the total number of neurons observed in unbiased snmC-seq2 profiling. This level of L5-ET neuron enrichment provides us with a unique opportunity to more closely investigate subpopulations of L5-ET neurons. In unsupervised clustering using genome-wide mCH levels measured in 100 kb genomic bins, L5-ET neurons further segregated into 15 subclusters upon uniform manifold approximation and projection (UMAP) embedding (Fig. 3a). Much of the separation between subclusters was driven by the source location of the neurons, as neurons from different source regions were clearly separated on the UMAP (Fig. 3b) and each of the subclusters consists of neurons mostly from one or two source regions (Extended Data Fig. 6a). In particular, RSP and AI each formed their own specific subcluster (cluster 13 and 3, respectively; Extended Data Fig. 6a, b). The similarities and differences between L5-ET neurons from different source regions were quantified using hierarchical clustering (Fig. 3c). The genome-wide mCH similarity is highest between MOp and SSp, followed by between VIS and AUD, and between PTLp and ACA. AI and RSP were more distinct; in particular, RSP was well separated from the remaining cortical regions. These similarities between source regions were not well explained by their spatial proximity anterior-posteriorly or medial-laterally, but better correlated with the anatomical and functional connectivity between these regions. For example, MOp and SSp are components of the somatic sensorimotor subnetwork, while AUD, VIS, ACA, and PTLp are components of the medial subnetwork that channels information between sensory areas (that include VISp and AUD) and higher order association areas (that include PTLp and ACA)^18^.

**Fig. 3.**
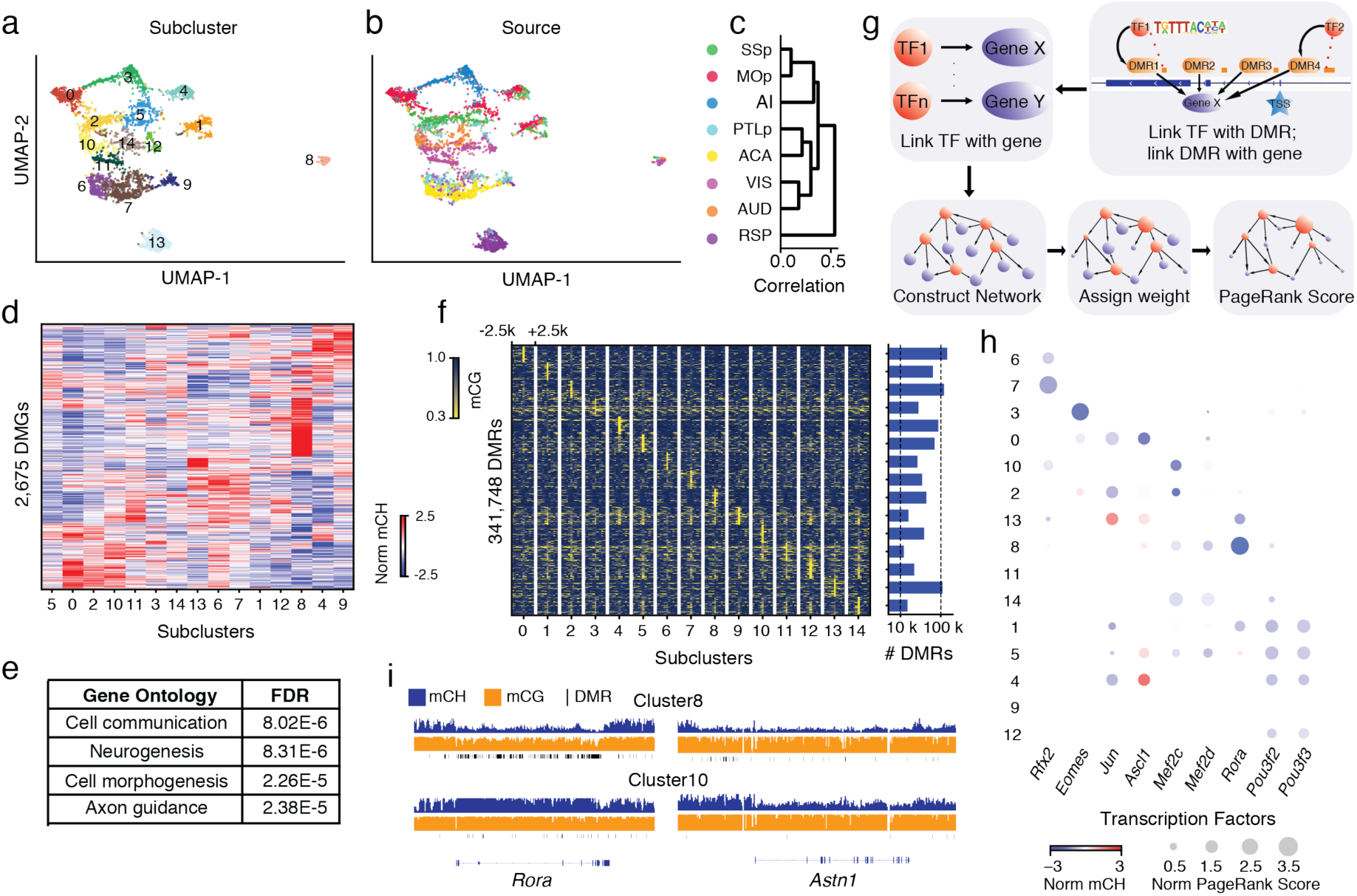
Epigenetic diversity of L5-ET neurons. **a, b**, Fifteen subclusters of L5-ET neurons (n=4,176) were identified and visualized on the uniform manifold approximation and projection (UMAP) plot generated using mCH levels in 100 kb genomic bins, colored by cluster (**a**), or the source region of neurons (**b**). **c**, Dendrogram shows the similarities between mCH profiles of L5-ET neurons from different source regions. **d, e**, In total, 2,675 CH-DMGs were identified in pairwise comparisons between L5-ET subclusters. Gene body mCH levels in each subcluster were visualized in the heatmap (**d**). Gene ontology (GO) enrichment of the CH-DMGs (**e**). **f**, Analysis of CG methylation (mCG) identified 341,748 differentially methylated regions (CG-DMRs) across the 15 L5-ET subclusters. The mCG levels at CG-DMRs and their 5kb flanking genomic regions in each subcluster were visualized in the heatmap (left). The numbers of CG-DMRs hypo-methylated in each subcluster were plotted in the bar chart (right). **g**, Workflow of the PageRank algorithm to infer crucial transcription factors. **h**, Examples of some predicted key regulator TFs are shown in the bubble plot. The size of each dot represents the normalized PageRank score of the TF. The color of the dot represents the gene body mCH of the TF in the corresponding L5-ET subcluster. **i**, Browser tracks of mCH (blue), mCG (orange), and CG-DMRs (black ticks) at *Rora* and its predicted gene target *Astn1*.

To further explore the molecular identity of these L5-ET subclusters, we used gene body mCH levels to identify cluster-specific genes. In total 2,675 CH-DMGs were identified in pairwise comparisons between subclusters (Fig. 3d, Supplementary Table 4; examples in Extended Data Fig. 6c), indicating that these genes have cluster-specific expression patterns. Gene ontology (GO) enrichment analysis revealed that these L5-ET subcluster CH-DMGs were enriched in genes involved in cell communication, neurogenesis, cell morphogenesis, and axon guidance (Fig. 3e, Supplementary Table 4).

In addition to identification of cluster-specific gene markers using gene body mCH, a powerful and unique advantage of methylation profiling is that cis-elements that regulate the marker genes can be predicted based on CG methylation. Differentially CG methylated regions (CG-DMRs) between clusters reliably mark cis-regulatory elements across the whole genome (not limited to gene bodies). Here, we identified 341,748 CG-DMRs that were hypo-methylated in the corresponding L5-ET subclusters (Fig. 3f, Supplementary Table 5). The average length of CG-DMRs was 227 bp, and 84.9% of them were distal elements that located more than 5kb from the annotated transcription start sites (TSSs).

The level of mCH at gene bodies is inversely correlated with gene expression, while the level of mCG at gene regulatory elements, such as promoters and enhancers, is inversely correlated with their regulatory activities. These relationships allowed us to use a gene regulatory network-based method to integrate this information and identify transcription factors (TFs) that might function as key regulators in each subcluster (see Methods; Fig. 3g). Specifically, in this network the nodes were genes (including TFs), while the edges connected the TFs to their potential target genes based on the TF binding motifs in CG-DMRs surrounding the TSSs. The weights of the nodes and edges were set according to the predicted expression levels (gene body mCH) of the genes. After applying a PageRank algorithm to score the genes in the network, we identified TFs that were potentially highly expressed and may regulate many other highly expressed genes in a subset of L5-ET clusters. This method combined the advantages of differential expression and motif enrichment analysis (Extended Data Fig. 6d, e), and enabled us to find TFs that may be expressed among a family of TFs sharing similar motifs^19^. For example, *Rora* (RAR Related Orphan Receptor A), a transcriptional activator, was scored as one of the top TFs and is hypo-CH-methylated in clusters 1, 8, and 13, and especially in cluster 8 (Fig. 3h, Extended Data Fig. 6d), indicating its potential expression. The binding motif of RORA was also enriched in the CG-DMRs of these same clusters, suggesting that RORA may bind to cis-regulatory elements that in turn regulate a set of predicted downstream target genes. Many of these target genes are related to brain functions and also hypo-methylated in cluster 8 (Extended Data Fig. 6f). For example, one of its predicted downstream target genes, *Astn1* (Astrotactin 1) is also hypo-CH-methylated in cluster 8 and encodes for a neuronal adhesion molecule, showing clear correlation between *Rora* and *Astn1* expression inferred from gene-body mCH (Fig. 3i).

### Subclusters of L5-ET neurons project to different targets

Our analyses of cortical IT neurons revealed epigenetic differences between neurons that related to both their cortical locations and their projection targets. Although the separation of L5-ET neuron subclusters was mostly driven by the source regions, neurons from the same source regions (except AI and RSP) distributed into more than one subcluster (Fig. 3a, b Extended Data Fig. 6b), prompting us to ask whether some of the differences between L5-ET subclusters also correspond to the different projection targets. To investigate this, we performed another iteration of clustering analysis using L5-ET cell data from each of the source regions separately, and identified finer L5-ET subclusters within each source region (Extended Data Fig. 7a). Consistent with these subclusters being related to true differences between putative cell types, all pairs of subclusters had more than 5 differentially CH-methylated 100 kb bins (CH-DMBs) (298 CH-DMBs on average).

We then examined whether neurons projecting to a specific target region were enriched or depleted in any of the subclusters (Extended Data Fig. 7c, d). Among all comparisons between projection targets and subclusters, neurons projecting to medulla (MY) were most distinct. SSp L5-ET neurons further segregated into seven subclusters (Fig. 4a), among which SSp→MY neurons showed a clear enrichment in subcluster 0 (FDR = 1.72E-2, Wald test; Fig. 4b, c). Similarly, we identified seven subclusters of MOp L5-ET neurons, and MOp→MY neurons were also significantly enriched in one of the subclusters (FDR = 6.81E-3, Wald test; Extended Data Fig. 7c, d). Moreover, MY-projecting neurons were robustly distinguished from other L5-ET neurons in our prediction models for both MOp and SSp (average AUROC = 0.929, 0.860; Fig. 4d, Extended Data Fig. 8a). Together, these analyses suggest that MY-projecting L5-ET neurons are more distinct than L5-ET neurons projecting to the other targets that were assessed.

**Fig. 4.**
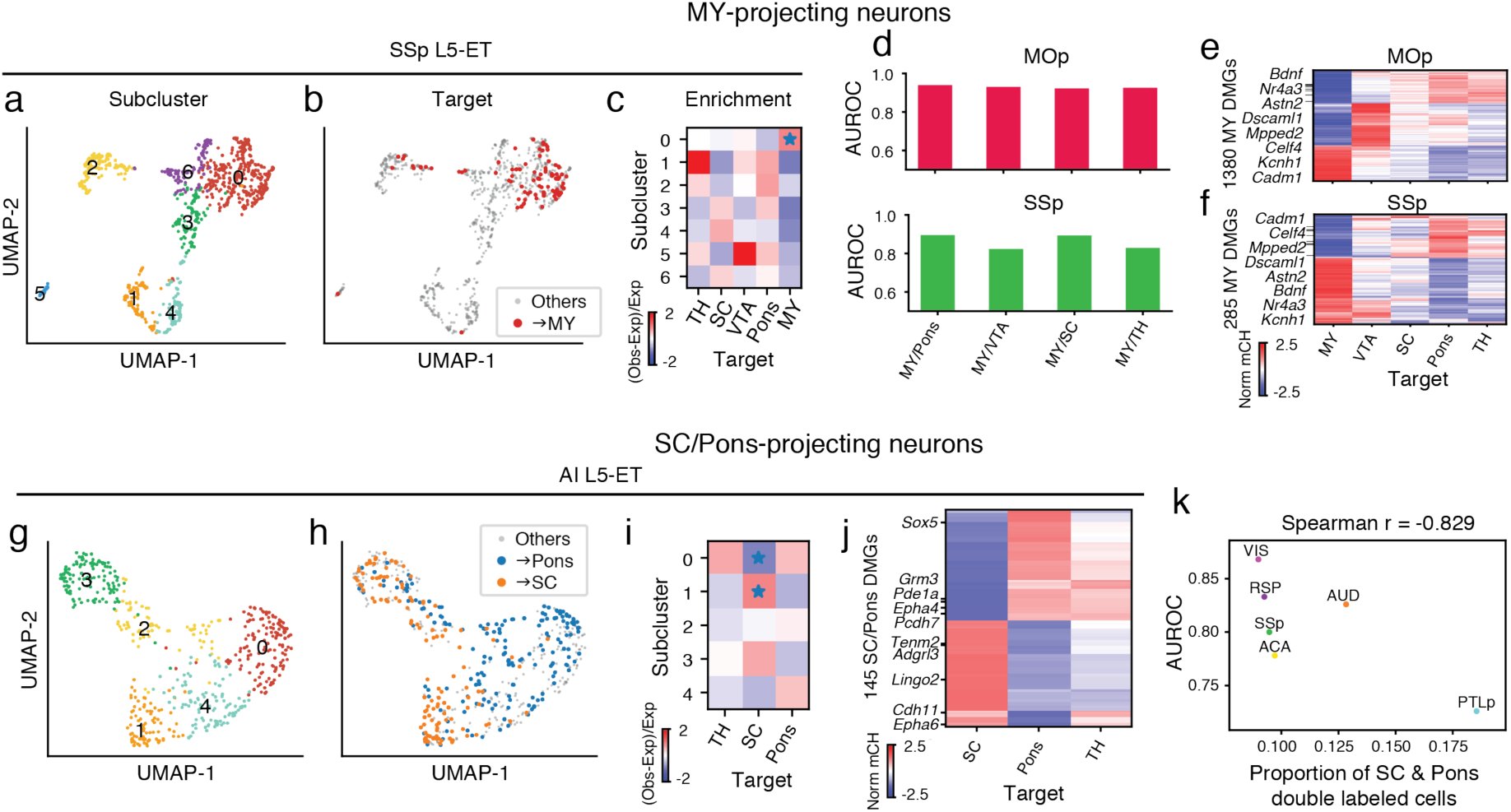
Epigenetic differences between L5-ET neurons projecting to different targets. **a-f**, L5-ET neurons projecting to MY had more distinct DNA methylation profiles than other L5-ET neurons: SSp L5-ET neurons (n=884) segregated into 7 subclusters as visualized on the UMAP plot generated using mCH levels in 100 kb genomic bins (**a**). Compared to other SSp L5-ET neurons, SSp→MY neurons occupied a distinct space on the UMAP that corresponded to SSp subcluster 0 (**b**). The enrichment of SSp→MY neurons in SSp subclusters was calculated and visualized in the heatmap (**c**; * represents FDR<0.05). We constructed prediction models to distinguish →MY neurons from →Pons, →VTA, →SC, and →TH neurons. AUROC scores showed that the models performed well in both MOp (**d**, top) and SSp (**d**, bottom) for comparisons between →MY neurons versus neurons projecting to each of the other targets. **e, f**, In total 1,380 CH-DMGs were identified in pairwise comparisons between MOp→MY neurons and MOp neurons projecting to another subcortical ET target. The gene body mCH levels of these CH-DMGs in MOp neurons projecting to each ET target were visualized in the heatmap (**e**). Similarly, 285 SSp→MY CH-DMGs were identified and plotted in the heatmap (**f**). Gene names for example CH-DMGs that were hypo-methylated in both MOp→MY and SSp→MY neurons are highlighted in the heatmaps (**e, f**). **g-k**, Epigenetic differences between Pons-projecting versus SC-projecting neurons varied across cortical regions: In AI, L5-ET neurons (n=531) separated into 5 subclusters as visualized on the UMAP plot (**g**). AI→Pons and AI→SC neurons occupied different positions on the UMAP (**h**), corresponding to their differential enrichment in AI subclusters 0 and 1 (**i**; * indicating FDR<0.05). 145 CH-DMGs were identified between AI→SC versus AI→Pons neurons. mCH levels of these SC/Pons CH-DMGs in AI→SC, →Pons, and →TH neurons were plotted in the heatmap (**j**). **k**, The variation of AUROC from prediction models to distinguish →SC versus →Pons neurons from different source regions suggested that the levels of distinction between →SC and →Pons neurons vary between cortical regions. From this observation, we hypothesized that different cortical regions had different proportions of neurons that made dual projections to both SC and Pons. The proportion of double labeled cells was negatively correlated with the AUROC score in each source area, supporting the hypothesis.

To investigate which genes drive the observed epigenomic differences between MY-projecting L5-ET neurons and other L5-ET neurons, we compared the gene body CH methylation profiles of MY-projecting L5-ET neurons to L5-ET neurons projecting to each of the other ET targets. In total, we identified 1,380 CH-DMGs between MOp→MY L5-ET neurons and at least one of the other ET projections (Fig. 4e, Supplementary Table 6). The majority of CH-DMGs were shared across the other ET projections. Specifically, among the 939 CH-DMGs that were hypo-methylated in MY-projecting neurons, 98 (10.4%) were universally hyper-methylated in all the other ET projections; Among the 441 CH-DMGs that were hyper-methylated in MY-projecting neurons, 85 (19.3%) were hypo-methylated in all the other ET projections. These results suggest that there are shared molecular differences that distinguish MOp→MY neurons from MOp neurons that project to VTA, SC, Pons, or TH. Similarly, 285 CH-DMGs were identified between SSp→MY L5-ET neurons and at least one of the other ET projections (Fig. 4f, Supplementary Table 6), among them 111 were hypo-methylated in SSp→MY neurons and 174 were hyper-methylated.

In total, 171 CH-DMGs were identified in both MOp→MY and SSp→MY neurons (a few examples highlighted in Fig. 4e, f), suggesting a general regulatory mechanism that may be shared by different cortical regions. Accordingly, models trained in either MOp or SSp to distinguish MY-projecting neurons usually performed well when tested in the other region (Extended Data Fig. 8b). Indeed, similar enrichment of MY-projecting neurons in subpopulations of L5-ET neurons has been reported in ALM using scRNA-seq (retro-seq)^13^. To compare these observations, we used gene body mCH as a proxy for gene expression to integrate our L5-ET Epi-Retro-Seq data with the ALM retro-seq data. Joint t-SNE showed that the MY-projecting L5-ET neurons were enriched in the same subcluster (Extended Data Fig. 9). *Slco2a1*, a marker gene of the ALM MY-projecting cluster^9,13^ is hypo-methylated in MOp→MY but not in SSp→MY neurons (Extended Data Fig. 9h). We identified *Astn2* as a marker gene for the MY-projecting L5-ET cluster in both MOp and SSp (Extended Data Fig. 9i). ASTN2 mediates the recycling of neuronal cell adhesion molecule ASTN1 in migrating neurons, and its deletion has been associated with schizophrenia. This suggests that, compared to other L5-ET neurons, MY-projecting neurons have distinct molecular properties, and these distinctions are likely shared across several cortical regions.

In addition to the MY-projecting L5-ET neurons, we also observed differences in genome-wide mCH profiles between other ET projections. For example, L5-ET neurons in AI were segregated into five subclusters (Fig. 4g), and AI→Pons and AI→SC neurons were enriched in different subclusters (Fig. 4h, i, Extended Data Fig. 8c). In contrast, AI→Pons and AI→TH neurons were enriched in similar subclusters (Extended Data Fig. 8c). Analysis of gene body mCH identified 145 CH-DMGs that were differentially methylated between AI→SC neurons versus AI→Pons, while most of them had similar expression patterns between AI→Pons and AI→TH neurons (Fig. 4j). Together, the results suggest that AI→Pons neurons are more distinct from AI→SC neurons and are similar to AI→TH neurons.

In contrast to the conservation across cortical areas ALM, MOp, and SSp for differences related to projections to MY, differences between Pons-projecting and SC-projecting neurons were not conserved across all cortical areas. We trained a prediction model using mCH profiles to distinguish Pons-versus SC-projecting neurons from different source regions. The model performed well in distinguishing the two projections from cortical regions AI (AUROC = 0.939) and VIS (AUROC = 0.868), but performed poorly in PTLp neurons (AUROC = 0.726) (Extended Data Fig. 8a). The AUROC scores were correlated with the counts of CH-DMGs identified between SC-projecting versus Pons-projecting neurons in the corresponding source regions (Spearman r=0.683). This suggests that the differences between Pons-projecting and SC-projecting neurons vary across the cortex.

From these observations, we hypothesized that the level of the epigenetic differences between the two projections might be correlated with the percentage of neurons that simultaneously project to both Pons and SC, which might vary between different cortical regions. That is, in a cortical area where more neurons project to both Pons and SC, the epigenetic profiles of Pons- and SC-projecting neurons might be expected to be less distinguishable in our data, and vice versa. To test this hypothesis, we performed double retrograde labeling of Pons and SC, and counted in each cortical source region the number of neurons labeled only by the tracer injected into Pons, only SC, or both (Supplementary Table 7). As our hypothesis predicted, PTLp had the highest percentage of double-labeled neurons, and in general the AUROC score from our model was negatively correlated with the percentage of double-labeled cells (Spearman r=-0.829, p=0.04) across the cortical regions (Fig. 4k). These correspondences are weak, however, for most source regions, so the correlation is driven primarily by the data from PTLp.

### L5-ET+CC neurons

Intriguingly, we noticed more than 30 VISp-projecting neurons in L5-ET clusters from ACA and RSP datasets (Fig. 5a, b). Since neurons in the L5-ET cluster are likely to project to ET targets, this finding suggested that some L5 neurons might project to both cortical and ET targets. These neurons were enriched specifically in one subcluster in ACA and RSP, respectively (FDR = 9.82E-5, 2.45E-3, Wald test; Fig. 5a-d). This type of subcluster in both RSP and ACA was marked by *Ubn2*, a highly expressed gene in visual systems, and many other genes also distinguished this cluster in either region.

**Fig. 5.**
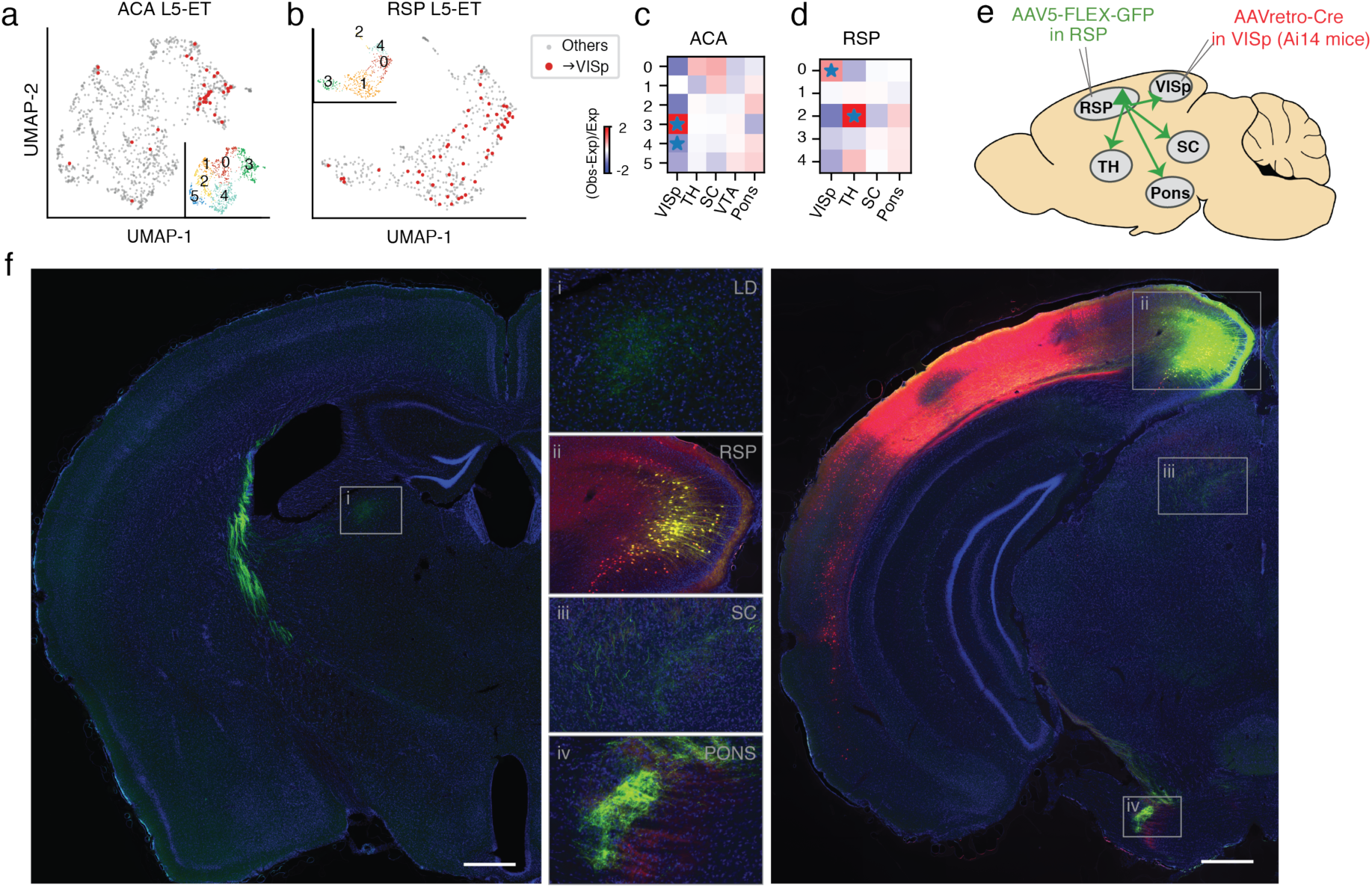
A L5-ET neuron type that projects to both ET and cortical targets (L5-ET+CC). **a**, UMAP embedding of ACA L5-ET neurons (n=1,131) using mCH in 100 kb bins, colored by projection targets (ACA→VISp in red, n=36) and subclusters (Inset). **b**, UMAP embedding of RSP L5-ET neurons (n=516) using mCH in 100 kb bins, colored by projection targets (RSP→VISp in red, n=53) and subclusters (Inset). **c-d**, ACA→VISp neurons were enriched in ACA L5-ET subcluster 3 and depleted from subcluster 4 (**c**). RSP→VISp neurons were enriched in RSP L5-ET subcluster 0 (**d**). (* indicating FDR<0.05). These observations suggested that some ACA and RSP neurons project to both ET and cortical targets (L5-ET+CC). To validate the existence of this L5-ET+CC cell type, we designed an anatomical labeling experiment as illustrated in **e**. AAVretro-Cre was injected into VISp of Ai14 (Cre-dependent TdTomato) mice, and AAV5-FLEX-GFP (Cre-dependent GFP) was injected in RSP. Therefore, RSP→VISp neurons, including their axonal projections, were selectively labeled with GFP. If RSP→VISp neurons also project to ET targets (L5-ET+CC neurons exist), GFP-labeled axons would be expected in subcortical ET targets such as SC, Pons, and TH. **f**, We performed these labeling experiments in three Ai14 mice and observed the same result in all mice. Examples of brain sections from one animal are shown. VISp neurons at the AAVretro-Cre injection site were labeled by tdTomato (red). RSP→VISp neurons were labeled with GFP (green), among which RSP→VISp neurons at the AAV5-FLEX-GFP injection site were labeled with both tdTomato and GFP (yellow; inset ii). Strong GFP signals of RSP→VISp axon terminals in subcortical ET regions were observed, including in the laterodorsal (LD) nucleus of the thalamus (inset i), SC (inset iii), and Pons (inset iv). Scale bars: 500 μm (low magnification).

Although, ET cells are generally thought to lack projections to other cortical areas, there is some evidence for such cells from previous studies^20^. Reconstructions of the axonal arbors of 24, L5 MOp neurons in rats revealed 3 neurons projecting to both SSp and TH^21^, and neurons in mouse secondary motor cortex have been shown to project to both AUD and ET targets^22^. In primates, single neurons projecting to both a cortical target, visual area MT, and a subcortical target, SC, have been observed in layer 6 of VISp^23,24^. However, since ET neurons represent a small percentage of primate neurons, these dual-projection neurons are extremely rare; they are also located in layer 6 rather than layer 5 making it difficult to predict whether they might be genetically more closely related to ET or to IT neurons, whether they might project to additional subcortical targets, or whether they might be unique to primates.

To anatomically validate our findings for RSP→VISp ET neurons in mice, we injected AAVretro-Cre in VISp and AAV-flex-GFP (Cre-dependent GFP) in RSP in three mice (Fig. 5e). This resulted in labeling of the complete axonal and dendritic arbors of RSP→VISp neurons such that their long-distance projections to locations other than VISp could be assessed. We observed strong GFP labeling of axon terminals in subcortical ET regions, including TH, SC, and Pons, in all three mice (Fig. 5f). These results indicate that single neurons in L5 of RSP can project simultaneously to both cortical and subcortical, ET targets in mice. Because these cells genetically cluster with L5-ET cells, we consider them a subtype of L5-ET cells that we refer to as L5-ET+CC. We do not use the term L5-ET+IT because many L5-ET neurons are known to project to another part of the telencephalon, the striatum.

## Discussion

Here, we have quantitatively analyzed and compared the methylation of mouse cortical neurons projecting to different cortical and subcortical target regions. We identified genes that were differentially methylated between different cortical areas projecting to the same targets, as well as between neurons in the same areas projecting to different targets. As expected from previous studies identifying IT- and ET-projecting neurons as distinct populations, these populations were also the most distinct in their gene methylation. We also identified differences between both IT neurons projecting to different cortical areas and between L5-ET neurons projecting to different ET targets. Cortical IT neurons projecting to different cortical targets were variable in the extent of their epigenetic differences. Some pairs of cortical target areas were more distinct than others and these epigenetic differences were often conserved across cortical sources areas. Differences between projection target pairs were typically larger than differences between cortical source areas for any given pair of projection targets.

Most distinct amongst the L5-ET neurons were those projecting to the medulla. This difference has been described previously for neurons in cortical area ALM^9^ and we find that this difference is conserved across the additional cortical areas that we analyzed, including MOp and SSp. In contrast, differences between L5-ET neurons projecting to SC versus pons were more distinct in some cortical areas (e.g. AI) than in others (e.g. PTLp). Dual retrograde tracer injections into both SC and pons revealed a corresponding difference in the proportions of double-labeled cells in different cortical areas, consistent with the expectation that neurons projecting to just one target can be different while those projecting to both targets cannot.

We found that a subpopulation of cortico-cortical RSP→VISp and ACA→VISp neurons clustered with L5-ET cells, contrary to the expectation that L5-ET and IT cortico-cortical cells are distinct populations. This suggested that some L5-ET cells might project to cortical targets and this hypothesis was validated anatomically. Our anatomical experiments showed that RSP→VISp cells do in fact project to many ET targets, including TH, SC and pons, and we refer to this cell type as L5 ET+CC. Although we found CC projection neurons that clustered with L5-ET cells for only two of the 26 CC projections that we sampled, there remain many other combinations that we did not test. Furthermore, previous studies have described L5 ET+CC cells in primary and secondary motor cortex^21,22^. It is therefore likely that future studies will reveal L5-ET+CC neurons in additional cortical areas projecting to various combinations of ET and cortical targets.

Finally, this large-scale effort linking methylation status directly to projection targets of mouse cortical neurons, allowed us to identify differences between projection cell types in TFs linked to differentially methylated regions. These observations provide insight into genetic mechanisms that might contribute to the differences in morphology and function of these cell types. As we have illustrated, this large dataset also provides the opportunity to predict regulatory elements that might be harnessed in future studies to target transgene expression to these cell types.

## Methods

### Experimental Animals

All experimental procedures using live animals were approved by the Salk Institute Animal Care and Use Committee. The knock-in mouse line, R26R-CAG-loxp-stop-loxp-Sun1-sfGFP-Myc (INTACT) was used for most experiments^4^ and they were maintained on a C57BL/6J background. 42-49 day old adult male and female INTACT mice were used for the retrograde labeling experiment. Adult C57BL/6J “wild-type” mice were used for double-retrograde labeling experiments.

### Surgical Procedures for Viral Vector and Tracer Injections

To label neurons projecting to regions of interest, injections of rAAV2-retro-Cre (produced by Salk Vector Core or Vigene, 2×10^12^ to 1×10^13^ viral genomes/ml, produced with capsid from Addgene plasmid #81070 packaging pAAV-EF1a-Cre from Addgene plasmid #55636) were made into both hemispheres of the INTACT mice. Animals were anesthetized with either ketamine/xylazine or isoflurane, placed in a stereotaxic frame, and 0.1 to 0.5 microliters of AAV was injected by pressure into stereotaxic coordinates corresponding to the desired projection target. A list of injection coordinates and volumes is provided in Supplementary Table 1. At least 2 male and 2 female mice were injected for each projection target. To label RSP neurons that project to VISp, RSP was injected with rAAV2-retro-Cre and VISp was injected with AAV-FLEX-GFP (Salk Vector Core) in each of 3 adult, Ai14 mice.

### Assessment of Double-Retrograde Labeling

To assess double-labeling of cortical cells projecting to Pons and/or Superior Colliculus, stereotaxic pressure injections of 0.1-0.2 microliters of 0.25-0.5% of Cholera Toxin Subunit B (CTB), Alexa Fluor 488 or 647 conjugated (Molecular Probes), were made into the pons and into SC of 4 mice. 6-7 days later, animals were perfused with phosphate buffered saline (PBS) followed by 4% paraformaldehyde in PBS. Brains were removed and sectioned coronally at 40 microns thickness with a freezing microtome. Sections were mounted and imaged with a 20X epifluorescence objective and images assessed to identify single and double-labeled neurons that were assigned to cortical areas. Only neurons in regions where labeled cells from both injections overlapped were counted. Therefore, some cortical areas in which there was no overlap are not included. For each animal, double labeled cells were quantified for each region as the proportion of double-labeled divided by the sum of all labeled cells. Mean values from the 4 animals are plotted in Fig. 4k.

### Brain dissection

Approximately two weeks after the AAVretro injection, brains were extracted from the 56-63 day old INTACT mice, immediately submerged in ice-cold slicing buffer (2.5mM KCl, 0.5mM CaCl_2_, 7mM MgCl_2_, 1.25mM NaH_2_PO4, 110mM sucrose, 10mM glucose and 25mM NaHCO_3_) that was bubbled with carbogen, and sliced into 0.6 mm coronal sections starting from the frontal pole. From each AAVretro-injected brain, the slices were kept in the ice-cold dissection buffer from which selected brain regions (Supplementary Table 1) were manually dissected under a fluorescent dissecting microscope (Olympus SZX16), following the Allen Mouse Common Coordinate Framework (CCF), Reference Atlas, Version 3 (2015) (Extended Data Fig. 1). The dissected brain tissues were transferred to prelabeled microcentrifuge tubes, immediately frozen in dry ice, and subsequently stored at −80°C.

### Nuclei preparation and single-nucleus isolation

For each dissected brain region, samples from 2 males and 2 females were pooled separately as biological replicates for nuclei preparation. The 2-mL glass tissue dounce homogenizer and pestles (Sigma-Aldrich D8938-1SET) were pre-chilled on ice. Nuclei were prepared using a modified protocol as reported by Lacar et al., 2016^25^. In summary, the frozen brain tissues were transferred to the dounce homogenizer with 1 mL ice-cold NIM buffer (0.25M sucrose, 25mM KCl, 5mM MgCl_2_, 10mM Tris-HCl (pH7.4), 1mM DTT (Sigma 646563), 10μl of protease inhibitor (Sigma P8340)), with 0.1% Triton X-100 and 5μM Hoechst 33342 (Invitrogen H3570), and gently homogenized on ice with the pestle 10-15 times. The homogenate was transferred to pre-chilled microcentrifuge tubes and centrifuged at 1000 rcf for 8 min at 4°C to pellet the nuclei. The pellet was resuspended in 1 mL ice-cold NIM buffer, and again centrifuged at 1000 rcf for 8 min at 4°C. The pellet was then resuspended in 450 μL of ice-cold NSB buffer (0.25M sucrose, 5mM MgCl_2_, 10mM Tris-HCl (pH7.4), 1mM DTT, 9ul of Protease inhibitor), and filtered through 40μM cell strainer. The filtered nuclei suspension was incubated on ice for at least 30 minutes with 50μl of nuclease-free BSA for at least 10 minutes, then incubated with GFP antibody, Alexa Fluor 488 (Invitrogen, A-21311) and anti-NeuN antibody (EMD Millipore MAB377) conjugated with Alexa Fluor 647 (Invitrogen A20173). GFP^+^/NeuN^+^ single nuclei were isolated using fluorescence-activated nuclei sorting (FANS) on a BD Influx sorter with 100μm nozzle, and sorted into 384-well plates preloaded with 2μl of digestion buffer for snmC-seq2^15^ (20 mL digestion buffer consists of 10 mL M-digestion buffer (2×, Zymo D5021-9), 1 ml Proteinase K (20 mg, Zymo D3001-2-20), 9 mL water, and 10 µL unmethylated lambda DNA (100 pg/µL, Promega, D1521)). The collected plates were incubated at 50°C for 20 minutes then stored at −20 °C.

### snmC-Seq2 library preparation

The bisulfite conversion and library preparation were performed following the detailed snmC-seq2 protocol as previously described^15^. The snmC-Seq2 libraries were sequenced on Illumina Novaseq 6000 using the S4 flow cell 2 × 150 bp mode.

### Reads processing and quality controls

We used the cemba-data pipeline to generate allc files from fastq files (<u>cemba-data.rtfd.io</u>), as described in Luo et al^6^. Specifically, the fastq files were first demultiplexed into single cells and trimmed of Illumina adaptors and 10 bp on both sides with Cutadapt^26^. The reads were mapped to mm10 INTACT mouse genome using Bismark^27^ with Bowtie2 aligner for each single end separately. The reads with MAPQ smaller than 10 were excluded. Potential PCR duplicates were removed with Picard MarkDuplicates. The reads from two ends were then merged to generate allc files using call_methylated_sites function in methylpy^28^. The global mCCC level was used to estimate the non-conversion rate of bisulfite treatment. The cells with less than 500 k non-clonal reads or non-conversion rate greater than 1% were removed from further analysis.

### Methylation data processing

For each single cell, we computed the methylated CH (*mc*) and total CH (*tc*) basecalls of all 100 kb bins across the genome and all gene bodies annotated in GENCODE vM10^29^. The autosomal bins that were covered by more than 100 basecalls in greater than 95% of cells were used for further analysis. The autosomal genes that were covered by more than 100 basecalls in greater than 80% of cells were used for further analysis.

### Computing posterior methylation levels

For each cell, we calculated the mean (*m*) and variance (*v*) of the mCH level across the 100 kb bins or genes. Then a beta distribution was fit for each cell *i*, where the parameters were then estimated by

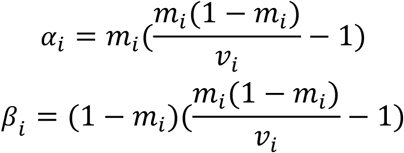

We then calculated the posterior mCH of each bin by

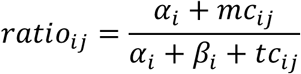

We normalized this rate by the cell’s global mean methylation by

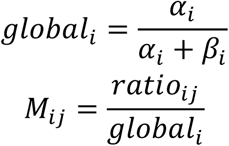

The values greater than 10 in *M* were set to 10. After normalization, *M*_*ij*_ is close to 1 when *tc*_*ij*_ is close to 0.

### Identification of highly variable bins

Highly variable methylation features were selected based on a modified version of the highly_variable_genes function in Scanpy^30^. In brief, since both the mean methylation level and the mean coverage of a feature (100 kb bin or gene) can impact methylation level dispersion^6^, we grouped features that fall into a combined bin of mean and coverage, and then normalized the dispersion within each group. After dispersion normalization, we selected the top 2,000 features based on normalized dispersion for dimension reduction.

### Removing potential doublets

By plotting all cells on t-SNE, we noticed a cell population that was located in the center of the plot and has a greater number of non-clonal reads than the others. To remove these potential doublets, we modified scrublet^31^ to adopt it to methylation data. Specifically, we first simulate the doublet cells by randomly selecting two cells in our dataset and sum the methylation/total basecalls of the two cells. Then the methylation levels of the simulated cells were computed using the posterior computing method. We simulated twice the number of doublets as the number of real cells. The top 2,000 highly variable features were selected for dimension reduction with principal component analysis (PCA) and the top 50 PCs were used to train a k-nearest neighbor (kNN) classifier (k=50) to predict a doublet score for each cell. Based on the histogram of doublet scores of real and simulated doublet cells, the cells with doublet score higher than 0.1 were removed from further analysis. After removing the potential doublets, 13,414 cells were kept for further analysis.

### Cell clustering and annotation

After removing potential doublets, the top 2,000 highly variable features were selected for dimension reduction with PCA. The top 50 PCs were used for t-SNE visualization and construction of kNN graph (*G*) with Euclidean distance (k=25). We use *A* to represent the connectivity of *G*, where *A*_*ij*_ is 1 if node *j* is among the 25 nearest neighbors of node *i*, otherwise 0. The edge weights of *G* were assigned as the jaccard distance of the connectivity matrix *A*. We ran Louvain clustering (https://github.com/taynaud/python-louvain) with resolution 1.2 to partition the cells into 31 clusters and merged these clusters into major cell types based on known marker genes. The 11,827 cells within neuronal cell clusters were selected for further analysis.

### Neighbor enrichment score

The score was used to quantify the enrichment of cells that belong to the same category among the neighbors of each cell. A higher score represents the cells are more likely to form clusters with the cells belonging to the same category rather than in the other categories. The advantage of this score is that it only considers the local effect so that would remain high if the cells in a category form several different clusters that dissimilar with each other. The score was computed as follows. Euclidean distances between each pair of cells were computed using the first 50 PCs. For each cell, we found its 25 nearest neighbors in the same category, and 25*r* nearest neighbors from other categories, where *r* is the ratio between total number of cells in other categories and total number of cells in the same category. The area under the receiver operating characteristic (AUROC) using distances between the cell and these neighbor cells for distinguishing the categories were defined as the neighbor enrichment score of this cell. The methylation pattern of male and female mice are highly similar on autosome; therefore, the two genders were treated as replicates in the analyses.

### Pairwise prediction of the source and target regions

Based on the sources, and targets, the neurons could be separated into groups. Each group contains the neurons projecting from a specific source to a specific target. To test the similarity of two groups of cells based on DNA methylation, we trained logistic regression models to predict the group label of each cell. The posterior of 100 kb-bin or gene body mCH were used as features. We split the cells into training and testing sets based on the gender of the mice where the cell came from. The area under the receiver operating characteristic (AUROC) from cross-validation was used to measure the performance of the model. The higher AUROC represents better ability of the model to present the group label, which indicated the two groups had larger mCH differences and were more distinguishable.

When the groups being studied contained cells from different clusters (e.g. cortical projecting neurons in one source), we up-sampled the training set to make it better capture the group differences rather than the differences of cell distributions across clusters. For example, when comparing neurons projecting to two different cortical targets, the cluster composition differences could make the model over-weight the features marking different clusters. To get rid of this bias, we randomly repeated the neurons from the under-representing group and ensured the two groups had the sample number of training samples in each cluster. The models were then trained and tested in the same setting as mentioned above.

Several reasons could contribute to a low prediction performance. 1) Some neurons make projections to several targets simultaneously. These could result in the neurons being captured by multiple retrograde labeling experiments of different targets. It would be impossible to predict a single label with our pairwise models for this type of neuron. 2) Some neurons project to different target regions but have tiny epigenetic differences. 3) The epigenetic differences between neurons projecting to different targets varies across replicates. In this study, male and female mice were treated as biological replicates after removing sex chromosomes. Although methylation patterns of autosomes are similar, differences between genders might still exist. 4) The contamination levels of some projections are high, which make larger noise and hinder the models to capture real signals. 5) The sample sizes of some projections are small, which make the learning more challenging.

If the cross source/cluster predictions (described below) performed better than the within source/cluster models, we would suspect that shared differences between neurons projecting to different targets exist across sources/clusters, and the major reason for lower accuracies of within source/cluster models might be 4) or 5) described above. To systematically distinguish 1) to 3), other anatomic and genetic validation are still needed.

### Cross source prediction

The logistic regression models were trained to predict the projection targets in one source and tested in the other source. The training set and testing set came from mice of different genders. Specifically, the final AUROC were the average of AUROCs by training in male mice and testing in female mice and by training in female mice and testing in male mice. For cortical targets, we up-sampled the training set in the same way as the above section.

### Cross cluster prediction

This analysis was specifically for CC projection neurons to study whether the mCH differences between projection neurons were shared or distinct across clusters (layers). The logistic regression models were trained to predict the projection targets in one cluster and tested in the other cluster. The training set and testing set came from mice of different genders.

### Identification of differentially CH-methylated genes (CH-DMGs)

Wilcoxon rank-sum test and t test were widely used to identify differential genes in single-cell studies^30^, which consider each cell as an independent sample. However, the cells from the same replicate, individual, or batch would be more similar than the cells from different ones. Therefore, considering all cells as independent samples would overestimate the statistical power in single-cell data. To address this problem and take the replicate-level variation into consideration, we used a linear mixed model for the differential analysis and performed paired-wise comparisons between groups. The posterior mCH level of 12,261 autosomal genes after coverage filters were used for these analyses. The posterior gene-body mCH was used as dependent variables. Each individual mouse was considered as a random effect. The global mCH levels and the gender of the mice were considered as fixed effects. Other fixed effects were determined based on the comparison. Specifically,

For DMGs between L5-ET clusters:

Gene_mCH ∼ cluster + gender + global_mCH + (1 | mouse)

For DMGs between cortical targets in each source:

Gene_mCH ∼ target + cluster + gender + global_mCH + (1 | mouse)

For DMGs between ET targets in each source:

Gene_mCH ∼ target + gender + global_mCH + (1 | mouse)

Each gene was tested separately, and two-sided Wald test was performed to estimate the *P* value for the effect being tested. FDR was computed for each pair of groups with the Benjamini/Hochberg process. The fold-change of each gene was computed by the average mCH across cells in one group divided by the average mCH across cells in the other group, with pseudo-counts of 0.1. The criterions for significance when testing difference variables were distinct and shown as follows. For DMGs between L5-ET clusters: absolute log fold-change greater than log1.5 and FDR smaller than 0.01. For DMGs between IT targets or between ET targets in each source: absolute log fold-change greater than log 1.25 and FDR smaller than 0.01.

### Identification of differentially CG-methylated regions (CG-DMRs)

To identify DMRs, we merged the allc files of individual cells assigned to the same cluster to create a pseudo-bulk allc table for each cluster. Then we selected all the CG sites and combined the methylation on two DNA strands for each CpG site. We run methylpy^28^ DMRfind to identify the DMRs and require the DMRs to contain at least 2 differentially methylated CpG sites (DMS).

### Inference of crucial transcription factors (TF) with PageRank

The method was modified from Taiji^19^ to integrate the information of both gene body and regulatory regions. The 537 motifs in JASPAR 2018 non-redundant core vertebrate database^32^ were used for these analyses. We scanned each of the motifs against the mm10 INTACT mouse genome with ame^33^ and *P* value cutoff as 1e-4. The DMRs between clusters were expanded 100 bp on both sides, and the ones overlapping with motifs were assigned to the corresponding TF. The DMRs were also assigned to the potential genes they regulated using GREAT^34^. The TFs were then linked with the target genes based on these DMRs that links to both the upstream TFs and the downstream genes. A gene regulation network was constructed where the nodes represented the genes and edges represented the links between TF genes and target genes.

To assign weights to the edges and initiate the node importance, the normalized *n*_*cluster*_ × *n*_*gene*_ methylation matrix (*M*) were min-max normalized across clusters to 0-1 by

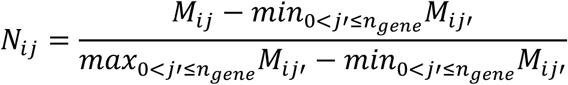

and 1 − *N*_*i*_ were used as the predicted expression of each gene in cluster *i*. The predicted expressions of all genes were used as starting importance *I*_0_. Then we used a *n*_*gene*_ × *n*_*gene*_ matrix *A* to represent the adjacency matrix of TF-gene regulation network, where *A*_*ij*_ was assigned as the predicted expression level of gene *i* if gene *i* is a TF. To ensure an undirected propagation, we used *B* = *A* + *A*^*T*^ as the final adjacency matrix. *B* was normalized by row into the transition matrix *P* by

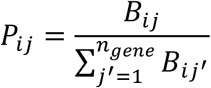

Next we performed a diffusion step of the PageRank scores through the network. For iteration *t*, the PageRank scores were computed by

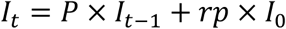

where *rp* represents a restart probability to balance the global and local effect of the propagation on the network. The diffusion step was stopped when |*I*_*t*_ − *I*_*t*_| < 10^−5^.

### Clustering of L5-ET cells in each source region

L5-ET neurons from Epi-Retro-Seq and unbiased snmC-Seq were combined in this analysis. After the same process as clustering all cells to derive posterior mCH level and select highly variable features, the first 30 PCs were used for computing kNN (k=15) and Louvain clustering. The resolutions used for source regions were 1.6 for MOp, AI, AUD, and RSP; 2.0 for SSp and PTLp; 1.0 for VISp; and 2.5 for ACA. The resolutions were determined based on visually examining the cluster numbers and projection enrichment.

To confirm that there were epigenetic features distinguishing the clusters, we computed the differentially methylated 100 kb bins (DMBs) across all pairs of subclusters using two-sided Wilcoxon rank-sum test. The bins were defined as differential if the absolute log fold-change between subclusters were greater than log 1.5, and FDR of the test smaller than 0.01. We also used AUROC>0.85 and AUPR>0.6 to define DMBs, which provided similar results. Two subclusters in RSP that had less than 5 DMBs were merged.

### Tests of projection enrichment in subclusters

As described above, the cells from the same replicate would be more similar, and considering all cells as independent samples will overestimate the statistical power in single-cell data. Therefore, we used linear mixed models to test for significant enrichment of particular projections in each subcluster, considering the mouse where the cells came from. The subclsuter was used as dependent variables. Each individual mouse was considered as a random effect. The projection target was considered as fixed effects. [Subcluster ∼ Target + (1 | mouse)]

Each projection target and each cluster were tested separately, and two-sided Wald test was performed to estimate the *P* value for the effect being tested. FDR was computed for each source with the Benjamini/Hochberg process. (Obs-Exp)/Exp in the enrichment matrices were computed using the same method as in Pearson’s chi-square test.

### Integration of Epi-Retro-Seq and Retro-Seq

Single-cell transcriptomic data from Tasic 2018^9,13^ was downloaded from NCBI Gene Expression Omnibus (GSE115746). 365 cells within clusters of ‘L5 PT ALM *Npsr1*’, ‘L5 PT ALM *Slco2a1*’, and ‘L5 PT ALM *Hpgd*’ were selected for integration analysis. The raw data was preprocessed using Scanpy^30^. Specifically, the read counts were normalized by the total read counts per cell and log transformed. Top 10,000 highly variable genes were identified and z-score scaled across all the cells. For methylation data, the posterior methylation levels of 12,261 genes in the 4,176 L5-ET cells were z-score scaled across all the cells and used for integration. We used Scanorama^35^ to integrate the z-scored expression matrix and minus z-scored methylation matrix with sigma equal to 100.

### Overlap score

Overlap score quantifies the similarity of the distributions of two groups of cells across clusters, where higher scores represent the two groups are more likely to be co-clustered. The scores were computed using the same method as in Hodge et al^14^. Specifically, a :*n*_*group*_ × *n*_*cluster*_ matrix *C* was first computed, where *C*_*ik*_ represents the number of group *i* cells in cluster *k*. *C* was normalized by row to *D*, and the overlap score between group *i* and group *j* was defined as 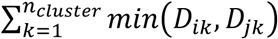.

## Data access and code availability

The data can be accessed via the NeMO ftp archive: http://data.nemoarchive.org/biccn/lab/callaway/projection/sncell/. The code for all of the analyses and the link to data browser can be found at https://github.com/zhoujt1994/Zhou2019.git

## Author contribution

Contribution to research design: E.M.C., Z.Z, M.M.B., J.R.E., J.Z., X.J., K.L.

Contribution to data collection: Z.Z., Y.P., A.R., E.W., C.L., M.A.K., A.F., P.A.M, A.B, A.A., M.V., L.B., C.F., J.R.N., R.G.C., M.R., M.J., T.I., B.D., J.B.S, C.O., M.M.B.

Contribution to data analysis: J.Z., P.T, Z.Z., E.M.C, M.A.K, A.F., H.L., S.N.

Contribution to data archive/infrastructure: E.A.M., Z.Z., Y.P., A.R., A.B.

Contribution to research coordination: Z.Z., E.M.C., J.R.E., M.M.B., Y.P., X.J., E.W., C.L., E.A.M., K.L.

Contribution to writing manuscript: J.Z., Z.Z., E.M.C., P.T., J.R.E., E.A.M., M.M.B.

## Acknowledgements

We thank Dr. Kai Zhang for advice on the PageRank algorithm, and Dr. Jesse R. Dixon for insightful comments. We are grateful to Dr. Michael Nunn for help with management of the project. This work is supported by NIMH U19MH114831 to E.M.C and J.R.E. M.A.K. is supported by NEI F31 EY028853. The Flow Cytometry Core Facility of the Salk Institute is supported by funding from NIH-NCI CCSG: P30 014195. J.R.E is an investigator of the Howard Hughes Medical Institute.

## Extended data figure legends

**Extended Data Fig. 1.**
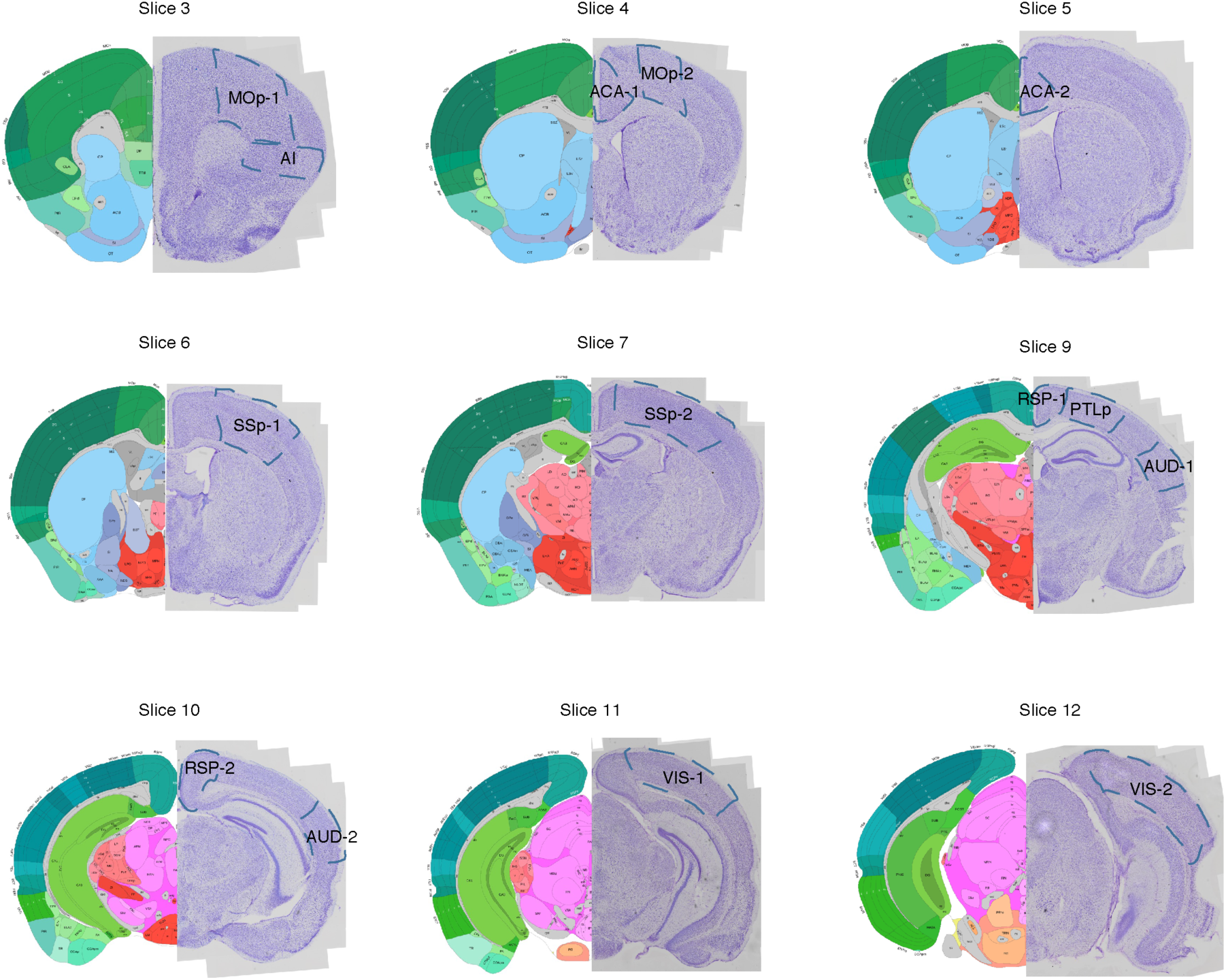
Source region dissection maps. The posterior views of dissected slices are shown. The slices correspond to Allen Reference Atlas level 33∼39 (slice 3), 39∼45 (slice 4), 45∼51 (slice 5), 51∼57 (slice 6), 57∼63 (slice 7), 69∼75 (slice 9), 75∼81 (slice 10), 81∼87 (slice 11), and 87∼93 (slice 12), respectively.

**Extended Data Fig. 2.**
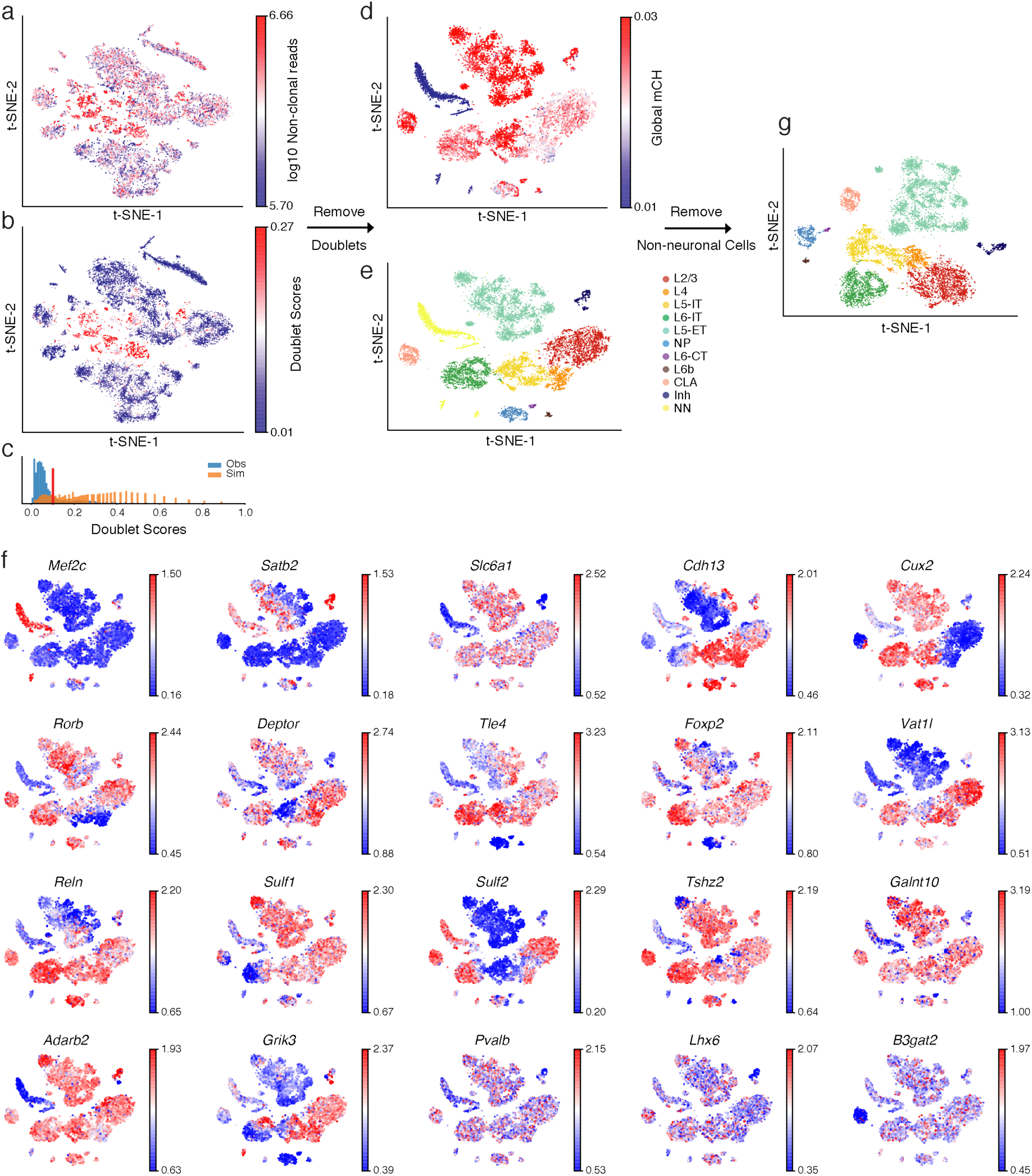
Removing potential doublets and non-neuronal cells. t-SNE of cells after quality control (n=16,971) colored by number of non-clonal reads (a) and predicted doublet scores (b). (c) Distribution of doublet scores for real cells (blue) and simulated doublets (orange). t-SNE of cells after removing doublets (n=13,414) colored by global mCH (d), cluster labels (e), and normalized gene-body mCH level of known cell type gene markers (f). Cells with low global mCH level are usually non-neuronal cells. t-SNE of single neurons (n=11,827) colored by the cluster labels (g). NN represents non-neuronal cells.

**Extended Data Fig. 3.**
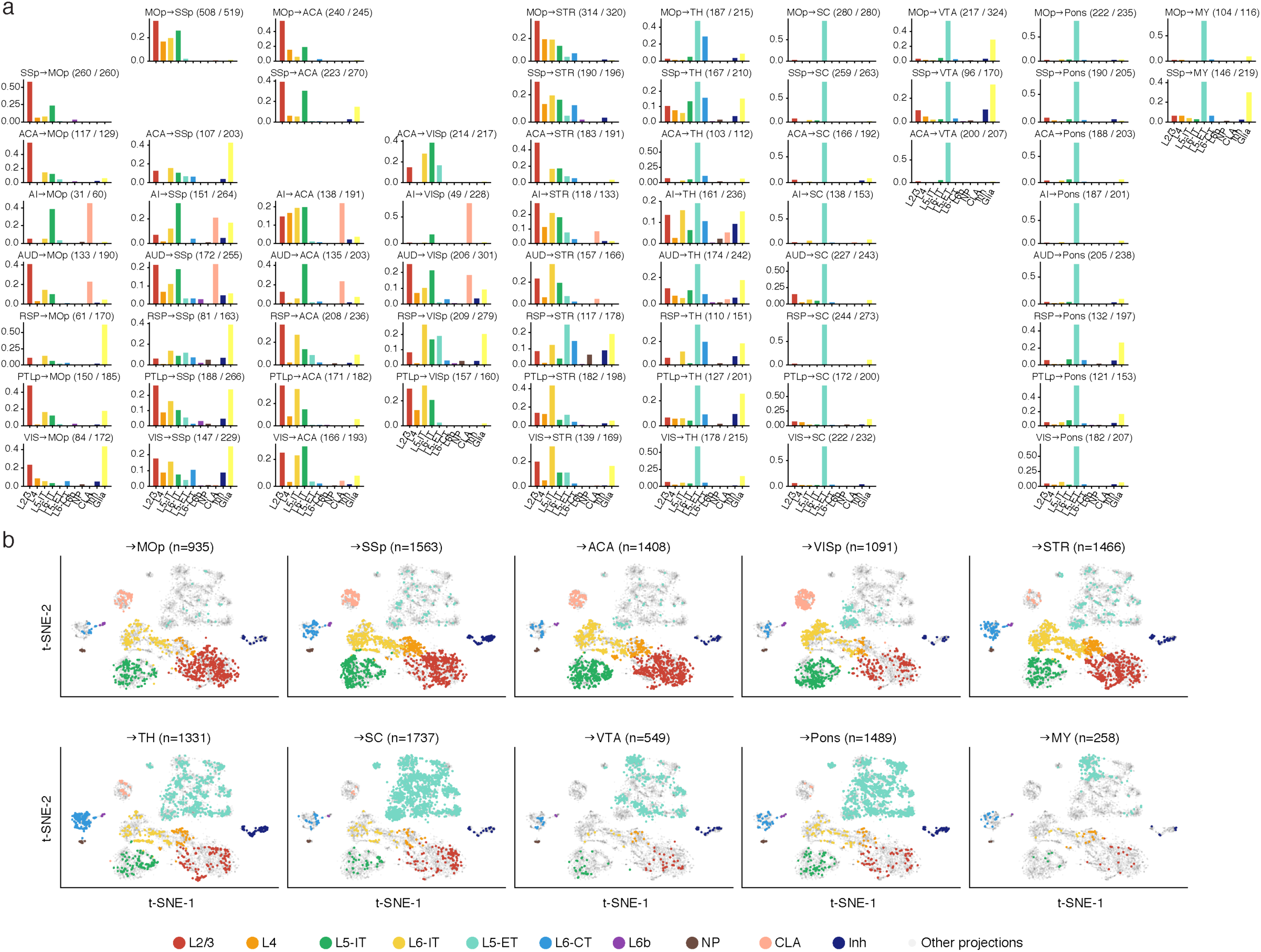
Cell type composition of all projections. (a) The proportion of cells projecting from each source region (row) to each target region (column) in all clusters including non-neuronal cells. (b) t-SNE of neurons (n=11,827) projecting to each IT target (top) and ET target (bottom). The cells projecting to the target were colored by clusters and cells projecting to all other targets were greyed.

**Extended Data Fig. 4.**
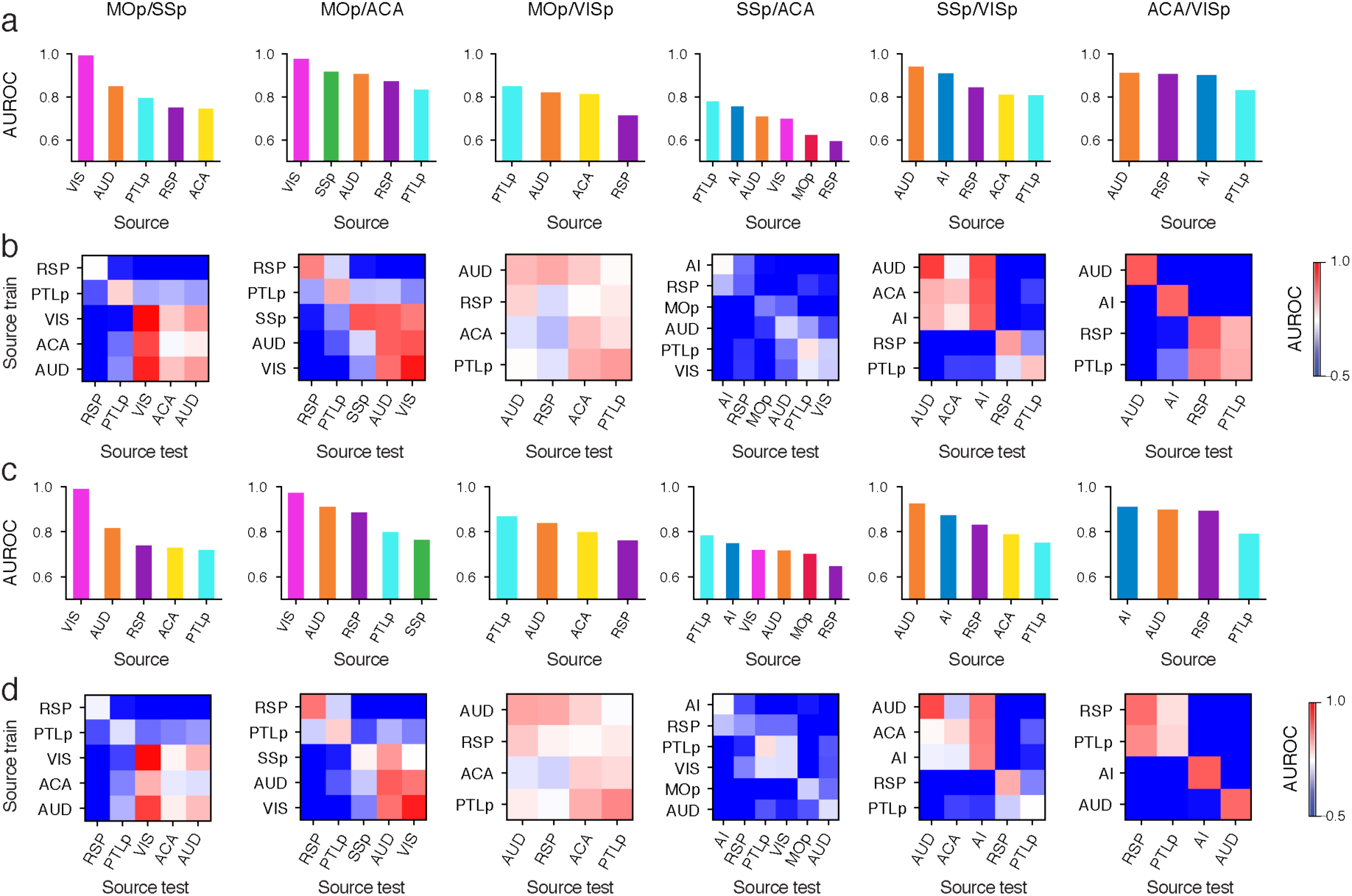
AUROC of cortical target pairs within and cross source regions. AUROC of models trained and tested in the same source region (a, c) or models tested in all source regions after trained in each one of them (b, d) using gene body (a, b) or 100 kb bin (c, d) mCH as features. The values in (a) and (c) correspond to the diagonals of (b) and (d) but ordered decreasingly.

**Extended Data Fig. 5.**
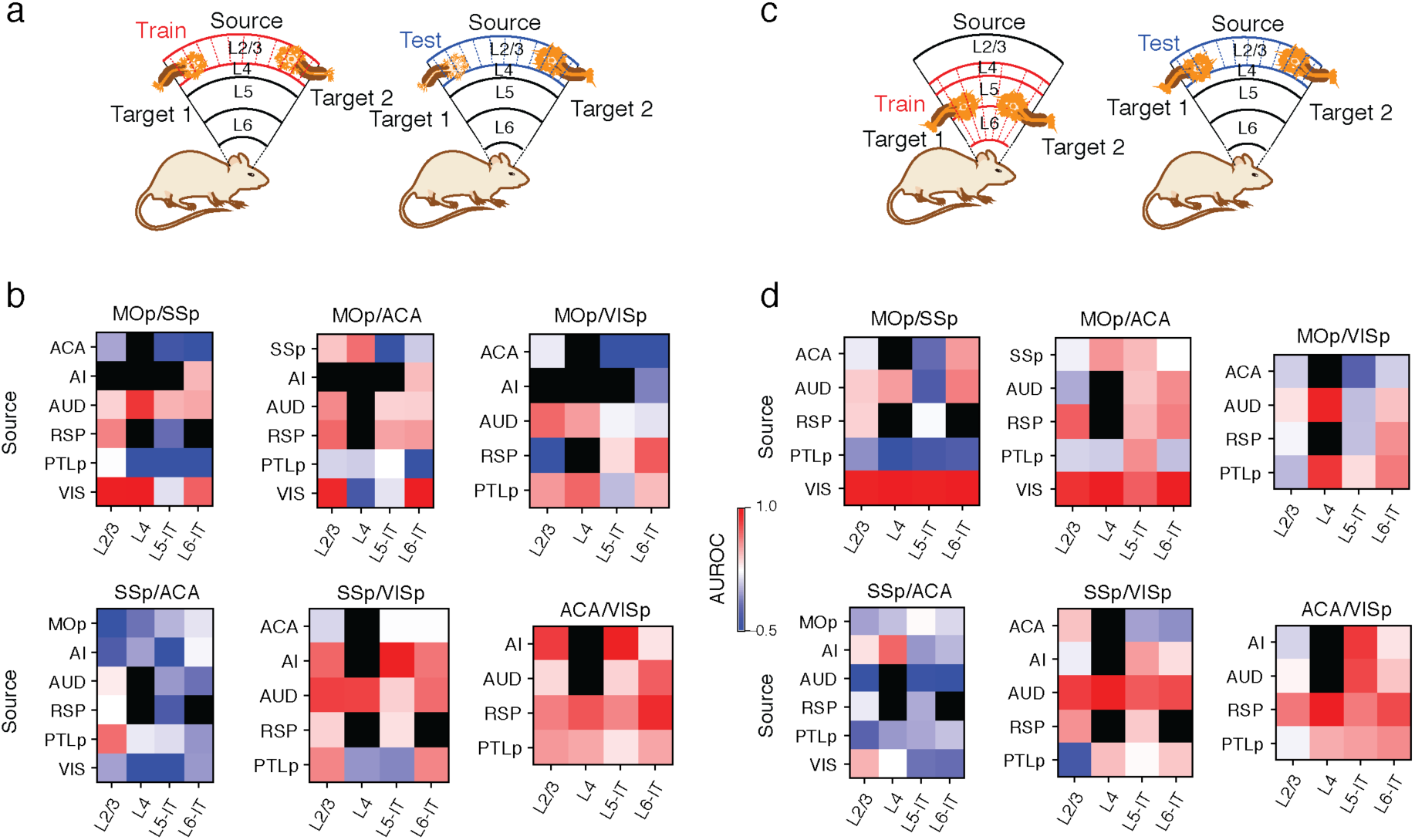
AUROC of cortical target pairs within and cross clusters. Demonstration of training and testing data for within layer prediction (a) and cross layer prediction (c). In (a), the models were trained and tested in the same layer with different replicates. In (c), the testing sets were the same as (a), but the models were trained in all other layers. AUROC of within layer prediction (b) or cross layer prediction (d). 100 kb-bin level mCH were used for all the predictions.

**Extended Data Fig. 6.**
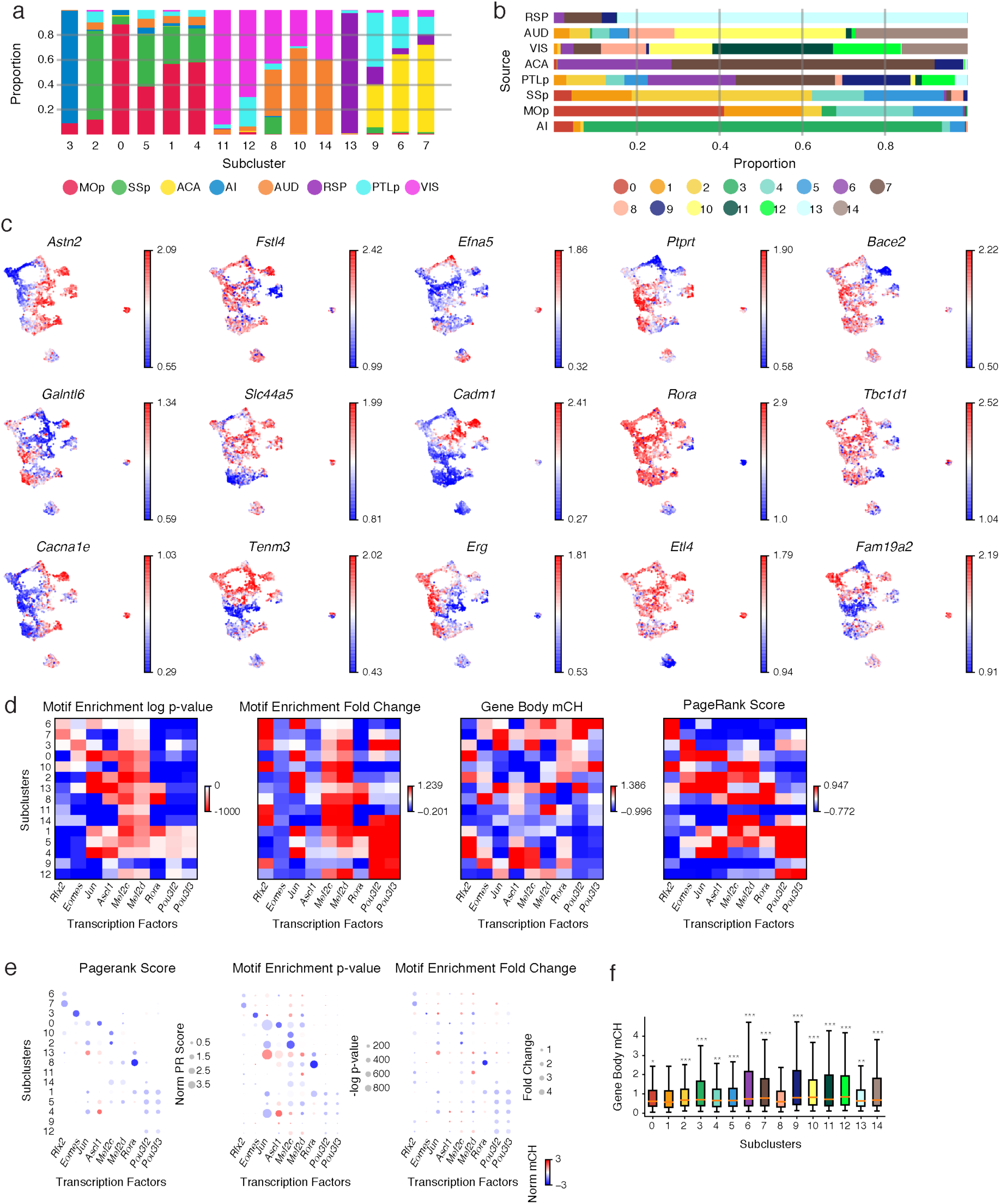
Signature genes and TFs of L5-ET subclusters. (a) Proportion of cells from all source regions in each subcluster. (b) Proportion of cells in all subclusters from each source region. (c) t-SNE of L5-ET cells (n=4,176) colored by the normalized gene-body mCH level of subcluster gene markers. (d) Motif fold-change within DMRs, and motif enrichment *P* value within DMRs, gene-body mCH, and PageRank score of the example TFs in all L5-ET subclusters. (e) Gene body mCH (color) against PageRank score (size, left), motif enrichment *P* value (size, middle), and motif enrichment fold-change (size, right) for the example TFs in all L5-ET subclusters. (f) Gene body mCH in all clusters of *Rora* target genes identified in cluster 8. Significances were determined by comparing cluster 8 with each of the other clusters (two-sided Wilcoxon rank-sum test). * represents p<1e-2, ** represents p<1e-3, *** represent p<1e-4. The elements of all box-plots are defined as: center line, median; box limits, first and third quartiles; whiskers, 1.5× interquartile range.

**Extended Data Fig. 7.**
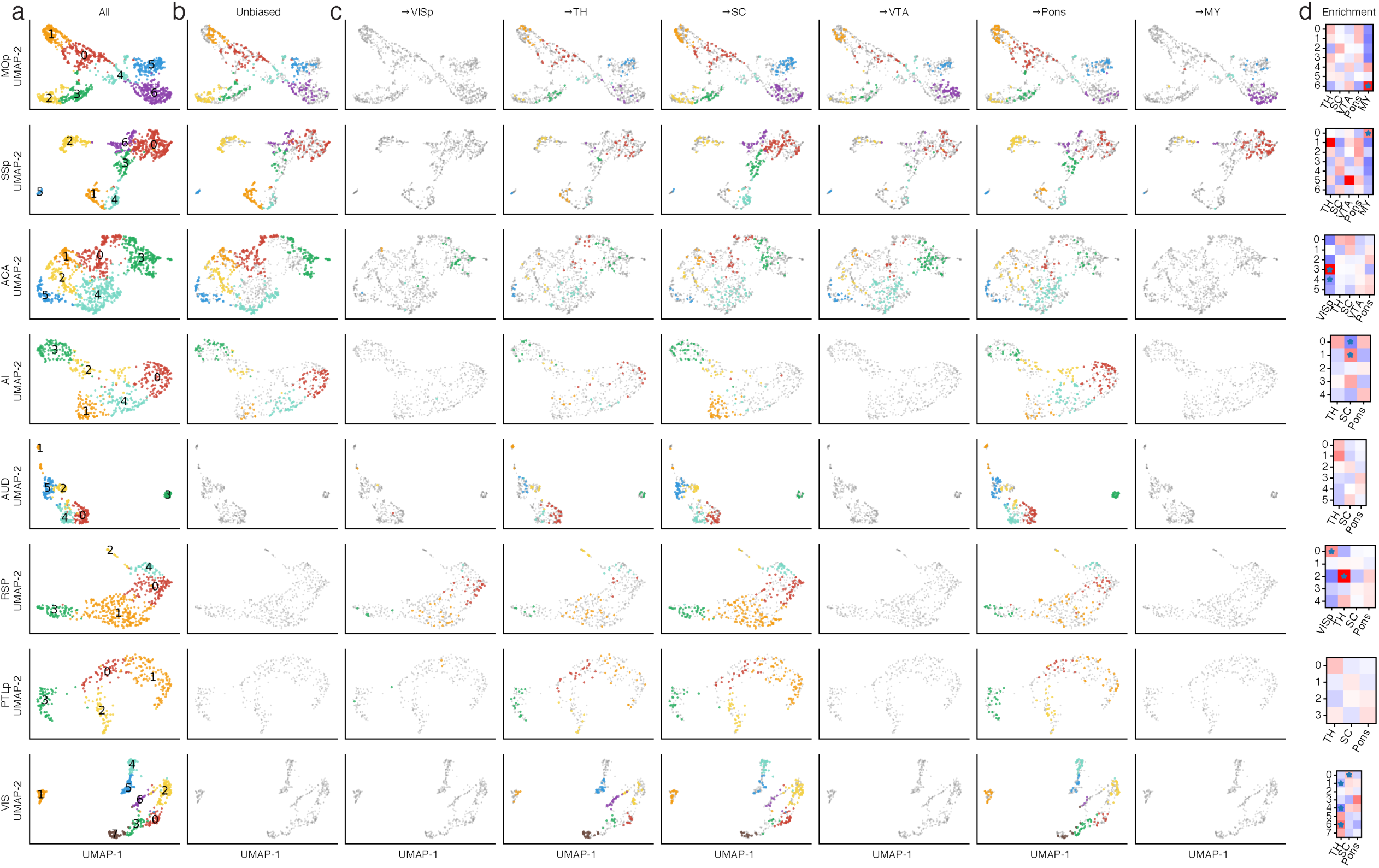
Enrichment of different projections in L5-ET subclusters. (a-c) t-SNE of L5-ET cells from each source region colored by subclusters. The colored cells are all cells (a), unbiased snmC-Seq cells (b), and cells projecting to each target (c). Other cells were greyed. (d) The enrichment of each projection in each L5-ET subcluster in each source. * represents FDR<0.05.

**Extended Data Fig. 8.**
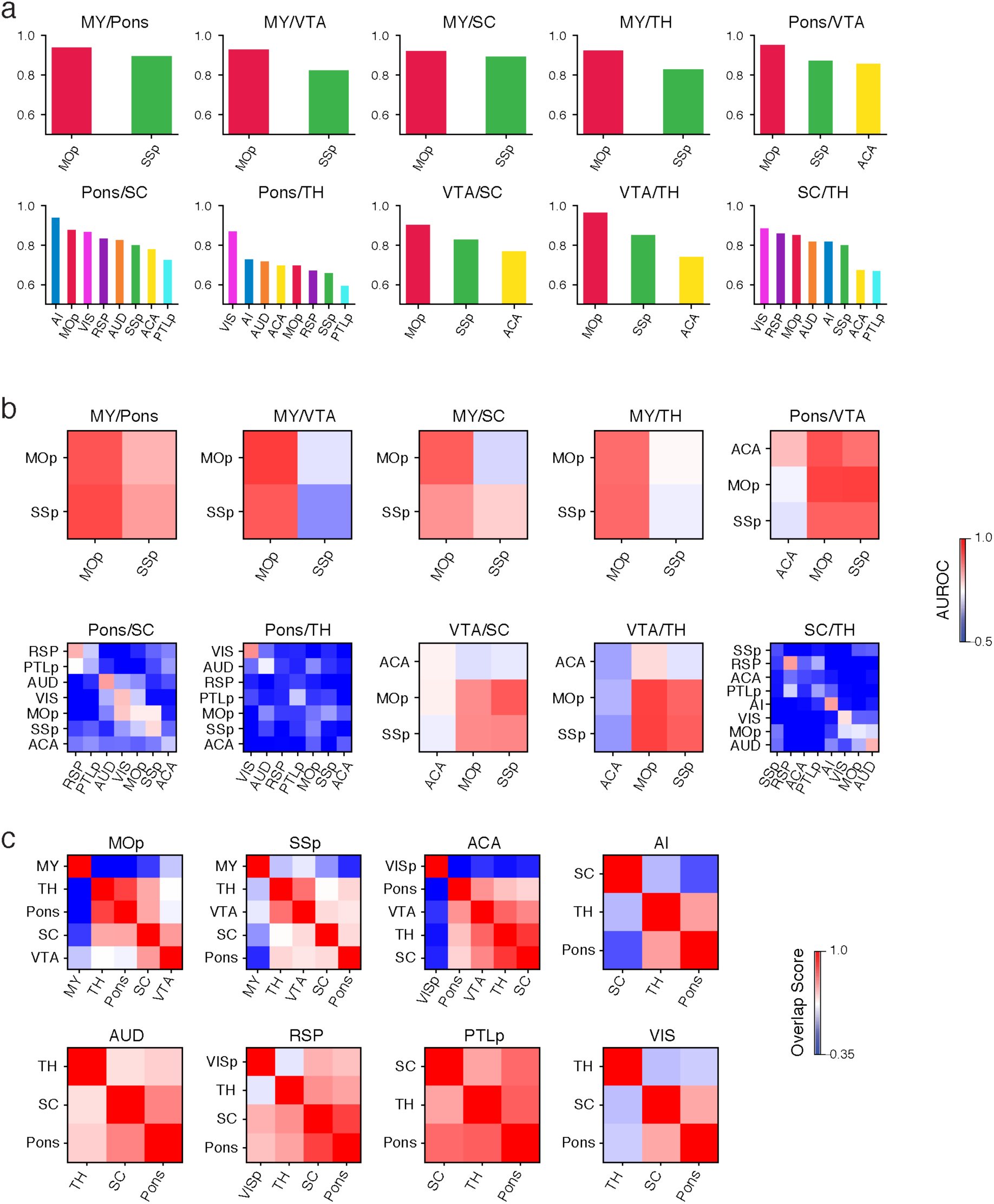
AUROC of ET target pairs within and cross source regions. AUROC of models trained and tested in the same source region (a) or models tested in all source regions after trained in each one of them (b) using 100 kb bin mCH as features. Training and testing sets were split by two-fold cross-validation in (a) to include AI, or split by replicates (b). (c) Overlap score between each pair of targets in each source region.

**Extended Data Fig. 9.**
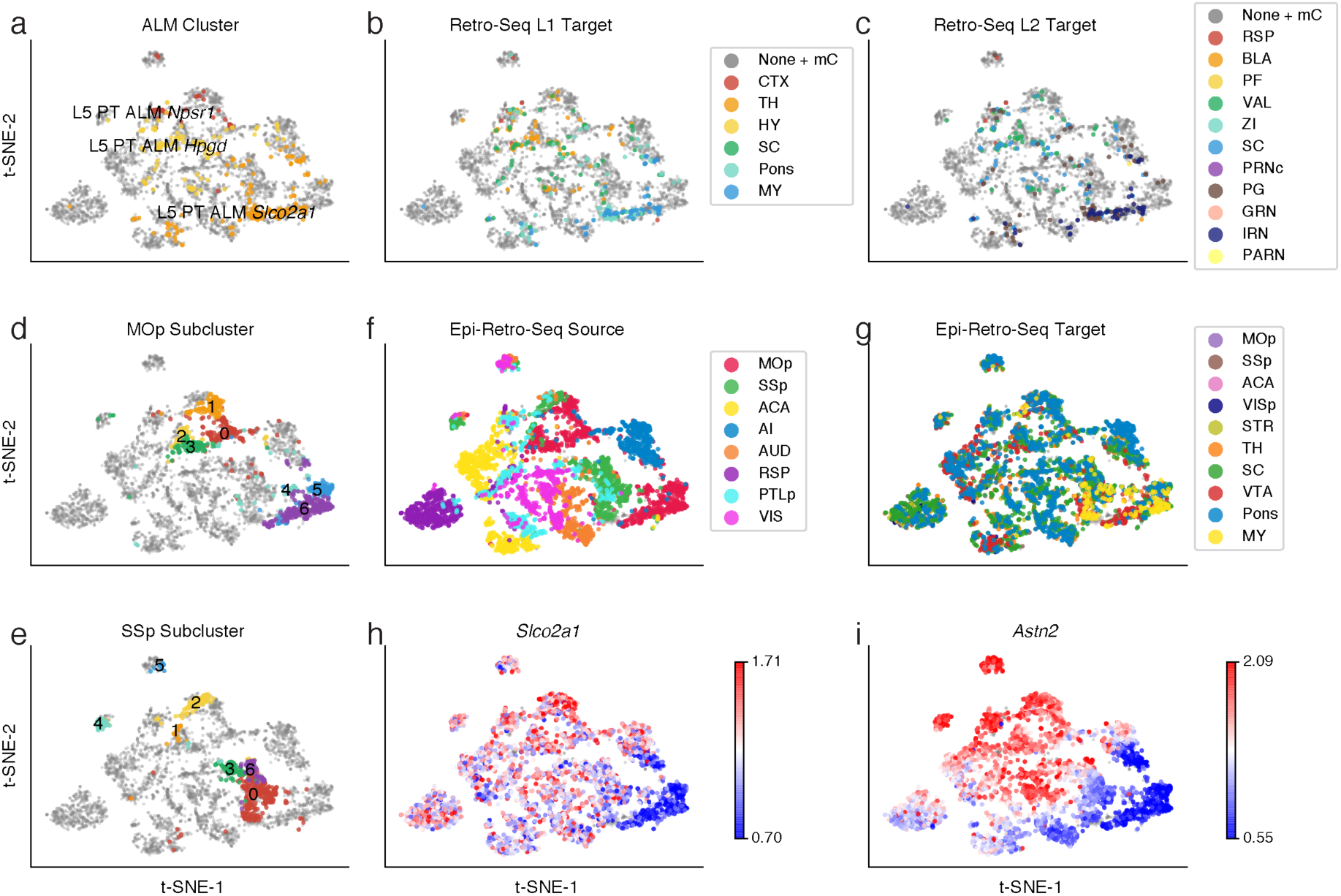
Integration of L5-ET cells from Epi-Retro-Seq and Epi-Seq. (a-c) L5-ET ALM cells in SMART-Seq (n=365) colored by clusters (a), major target regions (b), and detailed target regions (c). Epi-Retro-Seq cells were greyed. (d-i) L5-ET Epi-Retro-Seq cells from all source regions (n=4,176) colored by MOp subclusters (d), SSp subclusters (e), sources (f), targets (g), and gene body mCH of *Slco2a1* (h) and *Astn2* (i).

## Supplementary Tables

**Supplementary Table 1. Epi-Retro-Seq injection information**.

**Supplementary Table 2. Metadata and cluster assignment of 11**,**827 single neurons**.

**Supplementary Table 3. CH-DMGs between IT neurons projecting to different target regions and GO enrichment**.

**Supplementary Table 4. CH-DMGs between L5-ET subclusters and GO enrichment**.

**Supplementary Table 5. CG-DMRs between L5-ET subclusters and target genes assigned by GREAT**.

**Supplementary Table 6. CH-DMGs between L5-ET neurons projecting to different ET targets**.

**Supplementary Table 7. Cell counting in double labeling experiments**.

